# Rapid cognitive testing predicts real-world driving risk in commercial and medically at-risk drivers

**DOI:** 10.1101/2025.08.08.669388

**Authors:** Alice E. Atkin, Daniel Scott, Christopher Popowich, Anthony Singhal

## Abstract

Road safety is a major public and occupational health issue. Safe driving requires numerous cognitive and sensorimotor skills, and past literature suggests that cognitive testing can predict safe or unsafe driving in both healthy and medically at-risk drivers. However, such testing is often time-consuming and inaccessible. In this study, we designed a modified version of the Trail Making Test (TMT) which can be completed on a smartphone in approximately 5 minutes. We recruited 4405 commercially-licensed drivers and 314 medically at-risk drivers to complete the TMT, plus an on-road test of their driving abilities. We then trained and tested a logistic regression model using 50-50 splits on each dataset. The results of the model showed that the longer it took drivers in both groups to complete the TMT, the more likely they were to fail the on-road driving test. Accuracy for the commercial group was 83.8%, with a positive predictive value (PPV) of 34.4% and a negative predictive value (NPV) of 85.3%. Accuracy for the medically at-risk group was 63.1%, with a PPV of 55.8% and an NPV of 65.8%. Overall accuracy was 82.5%, with a PPV of 43.0% and an NPV of 84.3%. Log-transformed reaction time to targets was significantly associated with on-road failure in both driving groups. The results of this study suggest that a rapid and accessible version of the TMT can predict unsafe driving with comparable accuracy to more time-consuming and administratively burdensome means of testing.

## 1. Introduction

Road and traffic safety is a major global public health issue. According to the World Health Organization [WHO] in 2018, traffic fatalities were the leading cause of death in young people aged 5-29 worldwide, and the eighth leading cause of death amongst all ages. An estimated 1.35 million people die each year due to automobile crashes, with close to 50 million people being injured (James et al., 2020; WHO, 2018).

Safe driving requires interactions between multiple cognitive and sensorimotor functions such as, attention, memory, recognition, language, problem solving, and decision-making (Canadian Council of Motor Transport Administrators [CCMTA], 2022). Impairment to any of these functions can result in performance deficits, and an increased risk of crashes. Cognitive and sensorimotor impairment ranges in severity, and can be persistent or transient. Some of the primary etiology includes physical injury or disease, psychiatric illness, use of intoxicating substances, and situational factors such as active or passive fatigue (Hancock & Desmond, 2000; Saxby et al., 2013). Driver errors occur frequently when impaired by alcohol (Compton & Berning, 2015; Dingus et al., 2016) and/or cannabis (Bondallaz et al., 2016; Compton, 2017; Hartman & Huestis, 2013). All drivers are at risk of certain transient impairments, such as fatigue, but for the majority of impairment the risk is higher for certain drivers, such as repeat offenders who are responsible for the majority of impaired driving incidents (Clermont, 2018; Goldenbeld et al., 2016). Other causes of cognitive impairment, such as neurological disease, increase in prevalence over the lifespan (Dumurgier & Tzourio, 2020; Matthews et al., 2016). For example, mild cognitive impairment (MCI) can be due to neurological injury, but is also a precursor to dementia. The resulting impairment can often be treated with interventions and drug therapies, but many drivers with these conditions endure irreversible deterioration in their cognitive and sensorimotor skills until it is not safe for them to drive.

The significant dangers of impaired driving necessitates that in addition to administering tests when drivers are initially licensed, many jurisdictions have procedures for certain drivers to have their on-road driving skills re-tested as a condition of license renewal. Road tests are regularly required for drivers following license suspension due to drug or alcohol offenses, and medical experts can recommend that a patient take a road test whenever health-related impairment is suspected (CCMTA, 2022). While license renewal programs for older adults have the potential to identify cognitively impaired drivers and remove them from the road, the evidence of their effectiveness is minimal (Ichikawa et al., 2015; Langford et al., 2004; Mitchell, 2008; Rock, 1998; although see Vanlaar et al., 2016). It is certainly clear that impaired drivers should not be operating motor vehicles. Due to the uncertain effectiveness of age-based license renewal programs and their related administrative and cost burden, off-road cognitive and sensorimotor assessments are important tools for testing individual driver capability because they can signal that a subsequent on-road evaluation is needed. However, it is not always clear when an off-road assessment is required.

### 1.1 Driving Assessments

Individual assessments have historically been of value in occupational settings to determine job suitability (Goodstein & Lanyon, 1999; Kyllonen, 1986), particularly in work settings with higher levels of risk to safety, such as the military, and law enforcement (Hunt & Stevenson, 1946; Murphy, 1972). More recently, assessments have been used to evaluate an individual’s fitness for duty, which refers to their ability to perform a particular job without posing risk to themselves or others (Serra et al., 2007). Workplace safety incidents and injuries are often associated with cognitive errors, including memory failures and gaps in paying attention (Brossoit et al., 2019; Simpson et al., 2005; Wallace & Chen, 2005). This makes cognitive and sensorimotor assessments a valuable means for identifying performance risk and maximizing health and safety. Moreover, testing for impairment risk due to substance use in the transportation sector is critical, and needs further development for supporting road and driver safety in both the private and commercial sectors. (Pidd & Roche, 2014; Scott et al., 2023). To this point, one such test, the Cognitive Assessment Tool (DCAT) established in the 1990s by Al Dobbs and colleagues is based on a variety of tasks that measure cognitive ability and can be administered via computer in less than 30 minutes (Dobbs, 1997). The DCAT is currently used by healthcare systems and transportation companies, and performance on the assessment is predictive of whether drivers will subsequently pass or fail an on-road driving test (Dobbs, 1997; 2013; Dobbs et al., 1998; Korner-Bitensky & Sofer, 2009).

More recently, Impirica has developed a simpler to administer assessment named Vitals, which employs a handheld-tablet series of tests for simple reaction time, attention shifting, decision-making, memory, and sensorimotor control. In one study, our group showed that a population of drivers at-risk for cognitive impairment performed worse on the test battery relative to healthy controls, and the assessment results predicted on-road driving performance as measured by a trained instructor on city roads and highways (Bakhtiari et al., 2020). This assessment is sensitive to differences in impairment from cocaine, cannabis, or the combination of cocaine and cannabis (Tomczak et al., 2025). Importantly, the drug impaired groups also differed from adults without any drug impairment, and we assumed the impaired group members were unfit to drive. Two additional studies from our group examined the utility of Vitals in predicting on-road driving performance in a large sample of 3510 commercially licensed drivers for truck, bus, and light vehicles. In study 1, we showed with a logistic regression model that Vitals predicted on-road performance across all vehicle types, and that the drivers who failed the on-road evaluations demonstrated poorer performance on the judgement, memory, and sensorimotor aspects of the assessment (Scott et al. 2023). In study 2, we extended this work by replicating the results of Scott et al. (2023), with a new sample of data from 1924 drivers in the same commercial organizations (Atkin et al., 2024).

Taken together, these studies indicate that drivers who are likely to pose a considerable risk to themselves or others, and who are likely to fail an on-road driving test, can be reliably identified using an off-road, tablet-based screening tool that can be administered relatively quickly, efficiently, and at low cost. Indeed, the growing evidence strongly suggests that off-road impairment assessments can reliably predict risky driving across a range of populations and vehicle types (Atkin et al., 2024; Bakhtiari et al., 2020; Dobbs, 2013; Choi et al., 2015; Korner-Bitensky & Sofer, 2009; Scott et al., 2023).

### 1.2 Need for Rapid Assessment

In health care, reliable assessments are critical tools, and in the case of impairment disorders like dementia, timely assessments are crucial (O’Sullivan et al., 2005). In addition to health provider and care-giver/family observations, reliable assessments also employ tests of various cognitive factors. For example, as part of the National Dementia Strategy in the United Kingdom, there has been an emphasis of providing timely assessments to identify dementia risk as early as possible to improve treatment and support (Slater & Young, 2013). Such tests should be highly sensitive, fast to administer, easy to score, and efficient to infer the results. To this end, the most reliable tests like the MMSE and MoCA can be administered in less than 10 minutes, and the Abbreviated Mental Test (AMT), 6-item Cognitive Impairment Screen (6-CIT), Mini-cog, and the Clock Drawing Task (CDT) can be administered in less than 2 minutes (Slater & Young, 2013). The Sweet-16 (Fong et al., 2011) can be administered in less than 3 minutes and highly correlates with the MMSE. Importantly, some of the more rapid tests have been shown to reliably detect mild cognitive impairment (MCI). As a recent example, the Mini-cog has been shown to have 52% sensitivity and 80% specificity for MCI, and the CDT 56% sensitivity and 59% specificity (Tran et al., 2022).

While the medical community identifies the need for rapid cognitive screening tools for the prevention and treatment of disease, commercial industries with safety-sensitive operations like heavy trucking have been slower to identify the same need. Effective and reliable off-road cognitive assessments exist and have a key role to play, but the need for more rapid testing is as crucial in commercial environments as in clinical settings. Nearly 20% of road fatalities and traffic injuries in Canada between 2012 and 2021 involved commercial vehicles (Transport Canada, 2023). Many commercial operations run on tight delivery schedules where delays cost companies significant amounts of money (American Transportation Research Institute, 2024), and the drivers have capacity limits to their availability for work, and their ability to maintain appropriate levels of cognitive performance due to workload and fatigue (Bakhtiari et al., 2025). Thus, there is a clear need to develop reliable and more efficient tools to assess impairment for commercial settings. The question remains as to what kind of tests can be developed to be quick and efficient in terms of off-road administration, scoring, and interpretation, while maintaining industry standard reliability.

### 1.3 The Trail Making Test (TMT)

One cognitive test which has been frequently used to predict driving outcomes is the Trail-Making Test (TMT; Army Individual Test Battery, 1944; Reitan, 1958). The TMT has two levels of difficulty, TMT-A and TMT-B. TMT-A presents a single set of sequential items and requires people to connect them in sequence, while TMT-B presents two sets of sequential items and requires people to alternate between each set, item-by-item. TMT -A primarily tests visual search and processing speed, while TMT-B further tests executive functions such as cognitive flexibility and divided attention (Kortte et al., 2002; Salthouse, 2011; Sánchez-Cubillo et al., 2009). The tests are conventionally scored for completion time, although the number of errors can also be scored (Bowie & Harvey, 2006). Poor performance on the TMT is associated with traumatic brain injury (Lange et al., 2005; Periáñez et al., 2007), MCI and dementia (Ashendorf et al., 2008; Rasmusson et al., 1998; Venkatesan et al., 2018), and is also negatively correlated with education level and age in healthy adults (Kennedy, 1981; Mitrushina et al., 2005; Periáñez et al., 2007; Tombaugh, 2004). Originally a pencil-and-paper test, digital versions of the TMT have recently been created, with close correlations of scores across versions (Fellows et al., 2017), similar levels of clinical utility (Dahmen et al., 2017), and good construct validity and test-retest reliability (Zeng et al. 2017).

Due to the TMT’s extensive history as a neuropsychological test, many studies have explored the TMT’s potential as a tool for predicting unsafe driving. Because driving involves a variety of cognitive and sensorimotor functions, tests which assess these functions may have greater predictive value than a disease diagnosis or other physical indicators of health (CCMTA, 2025; Edwards et al., 2010). The TMT is advantageous for this purpose because of its short administration time (∼10 minutes) and ease of implementation.

Performance on TMT-B in particular is considered a marker of potential driving impairment, with the Canadian Medical Association’s <CMA> Driver’s Guide (2025) recommending that dementia patients undergo in-depth driving assessments if they score poorly on this test. TMT-B completion time has been found to be associated with driving performance in older drivers (Classen et al., 2008, 2013; Tarawneh et al., 1993), and in patients with Parkinson’s disease (Classen et al., 2015), stroke (Marshall et al., 2007; Motta et al., 2014), or cognitive impairment (Krasniuk et al., 2023; Ott et al., 2013). A study by Ball et al. (2006) found that TMT-B completion times are associated with future at-fault crashes in older drivers, as have retrospective studies by Stutts et al. (1998) and Friedman et al. (2013). Overall, a review of executive function tasks by Asimakopulos et al. (2012). A later study by Vaucher et al. (2014) concluded that TMT scores in older drivers lack the necessary sensitivity, specificity, and especially positive predictive value to recommend driving cessation on that basis alone, although poor scores were nevertheless associated with increased risk of poor on-road performance. The study used a lower-than-typical time cutoff (150s) for TMT-B failure, however, and using a strict criterion for identifying impaired cognition could have reduced the PPV by correctly identifying more unsafe drivers at the cost of more false alarms. A more recent review and meta-analysis by Rashid et al. (2020), which focused specifically on dementia patients, also found mixed results, with three out of five studies showing an association between TMT-B performance and driving. The preponderance of evidence nevertheless supports an association between TMT-B performance and driving.

TMT-A may also have utility for predicting driving outcomes. In a study of dementia patients by Carr et al. (2011), both TMT-A and TMT-B completion times differed significantly between drivers who passed and failed a road test, but their best predictive model only included TMT-A, not TMT-B. Likewise, Costello et al. (2024) found that TMT-A scores are a good predictor of on-road driving performance in older drivers, bettered only by the Useful Field of Vision (UFoV) test. TMT-A completion time was also associated with recent crashes in older drivers (Stutts et al., 1998). One advantage of TMT-A seems to be that it is easier, which means that very old and clinical populations can complete the test in the allotted time and without too many errors (Duncanson et al., 2018; Venkatesan et al., 2018). Duncanson et al. (2018) compared both older drivers and cognitively impaired drivers on both TMT-A and TMT-B, finding that TMT-A was discriminative of on-road driving performance in the patients while TMT-B was discriminative in healthy drivers. The authors point out that TMT-B involves cognitive functions such as set shifting which may not be essential for driving, so some patients who perform poorly on TMT-B due to their condition may nevertheless retain the necessary skills for safe driving.

The ability to predict safe driving in young and middle-aged adults with cognitive testing has been explored much less often, given the rarity of cognitive impairment amongst these populations. However, some such studies do exist. Kim et al. (2021) compared old and young (< 65) drivers on measures of simulated driving, using the TMT and other cognitive tests as predictors. The results showed that the TMT differed significantly between the young and old drivers and was highly discriminative of simulated driving performance. By contrast, Di Meco et al. (2021) found that TMT-A and TMT-B scores did not correlate with measures of driving performance, nor were they significant predictors of performance, although their sample only included teenagers and young adults (age 15-25), most of whom were women. Although middle-aged adults (age 25-64) are statistically the least likely age group to be involved in fatal crashes in most countries (International Transport Forum [ITF], 2024), they are also the most likely age group to be employed as drivers. If commercial drivers perform poorly on cognitive tests such as the TMT, this may predict unsafe driving due to occupational stress and the increased complexity involved in operating commercial vehicles, even if their cognitive performance does not fall to the level of clinical concern.

Some complicating factors when assessing the utility of the TMT for predicting driving include uncertainty around the optimal cut-off score for distinguishing good from poor performance (Duncanson et al., 2018; Roy & Molnar, 2013), and the fact that for TMT-B in particular, many older adults and clinical patients fail the test by taking too long or making too many errors (Frank et al., 2018; Papandonatos et al., 2015). Conventionally, patients are considered to fail TMT-B if they take more than 3 minutes to complete it, or make 3 or more errors, but some researchers have recommended shorter cutoffs for assessing driver fitness (Roy & Molnar, 2013; Vaucher et al., 2014) while others recommend that researchers use trichotomous cutoffs, independently validate cutoffs for the specific population they are working with, or do not use them (Classen et al., 2013; Molnar et al., 2009; Papandonatos et al., 2015).

### 1.4 Summary

Based on the foregoing review, it is established that the TMT has significant utility for predicting on-road driving performance and that such an assessment is important for road safety. Furthermore, there is a clear need for more rapid assessment tools that have the predictive value of other time-consuming and costly assessments. To address these two main points, we administered a rapid, smartphone-based cognitive assessment tool, called Neurapulse, to a large sample of mostly older, medically at-risk drivers, and commercial drivers. The assessment consisted of three tasks: a motor task, and both parts of the TMT. Using a similar approach to Scott et al. (2023) and Atkin et al. (2024), the drivers also completed an on-road driving evaluation, and we used a logistic regression model to predict the outcome of the on-road evaluation based on Neurapulse performance.

## 2. Methods

### 2.1 Participants

Our initial sample (n=5655) consisted of all available participant data, including medical and commercial sources, within our provided proprietary dataset. The target dataset consisted of a subset of participants that: A) fully completed an in-office battery that included a Motor task, a modified TMT-A task, and a modified TMT-B task under typical conditions, and B) fully completed a standardized road course under typical marking conditions. To achieve this target, we applied a set of exclusion criteria to filter out data collection that was incomplete (see Table 1) or poor quality. Several of the samples were excluded because they did not include any of the in-office tasks targeted for our analysis (n=109), while others were missing partial Motor task (n=9) or Trails task (n=17) data. If a data source, which included collection from various medical and commercial testing sources, did not use standardized marking (n=364) or had lower than 80% in-office/on-road test completion (n=434), all data from those sources were excluded. Three tests were excluded because the test was marked as compromised by the evaluator due to the participant not being able to complete it on their own. The full list of counts involved in each exclusion rule can be seen in Table 1.

**Table 1:** Participants affected by exclusion rules (n=936).

| Exclusion criterion | N |
| --- | --- |
| Missing all Trails/Motor task data | 109 |
| Incomplete Trails A/B task | 17 |
| Incomplete Motor task | 9 |
| Unstandardized marking data source (on-road course) | 364 |
| Incomplete in-office/on-road completion | 434 |
| Individual's test was marked as compromised on the participant record | 3 |
| **Total excluded** | **936** |

Table 2 contains the final participant counts after exclusions are applied. There were a total of 314 medically-sourced participants (176 safe, 138 unsafe) and 4405 commercially-sourced participants (3733 safe, 672 unsafe). Due to procedural constraints, we were unable to accurately record the gender of the participants. These constraints are shared with our previous work (Atkin et al., 2024; Scott et al., 2023). For a fuller explanation of the reasons for not measuring or estimating gender, see Atkin (2025, pg. 233-237).

**Table 2:** Safety group sample sizes for drivers of medical and commercial data sources in the full dataset (n=4719).

| Age Group | Safe Commercial | Unsafe Commercial | Safe Medical | Unsafe Medical | Total |
| --- | --- | --- | --- | --- | --- |
| <20 | 61 | 5 | 0 | 0 | 66 |
| 21-30 | 1140 | 215 | 3 | 1 | 1359 |
| 31-40 | 1099 | 203 | 3 | 0 | 1305 |
| 41-50 | 717 | 136 | 8 | 3 | 864 |
| 51-60 | 508 | 76 | 11 | 7 | 602 |
| 61-70 | 198 | 32 | 34 | 19 | 283 |
| 71-80 | 10 | 5 | 68 | 58 | 141 |
| 81+ | 0 | 0 | 49 | 50 | 99 |

**Figure 1:**
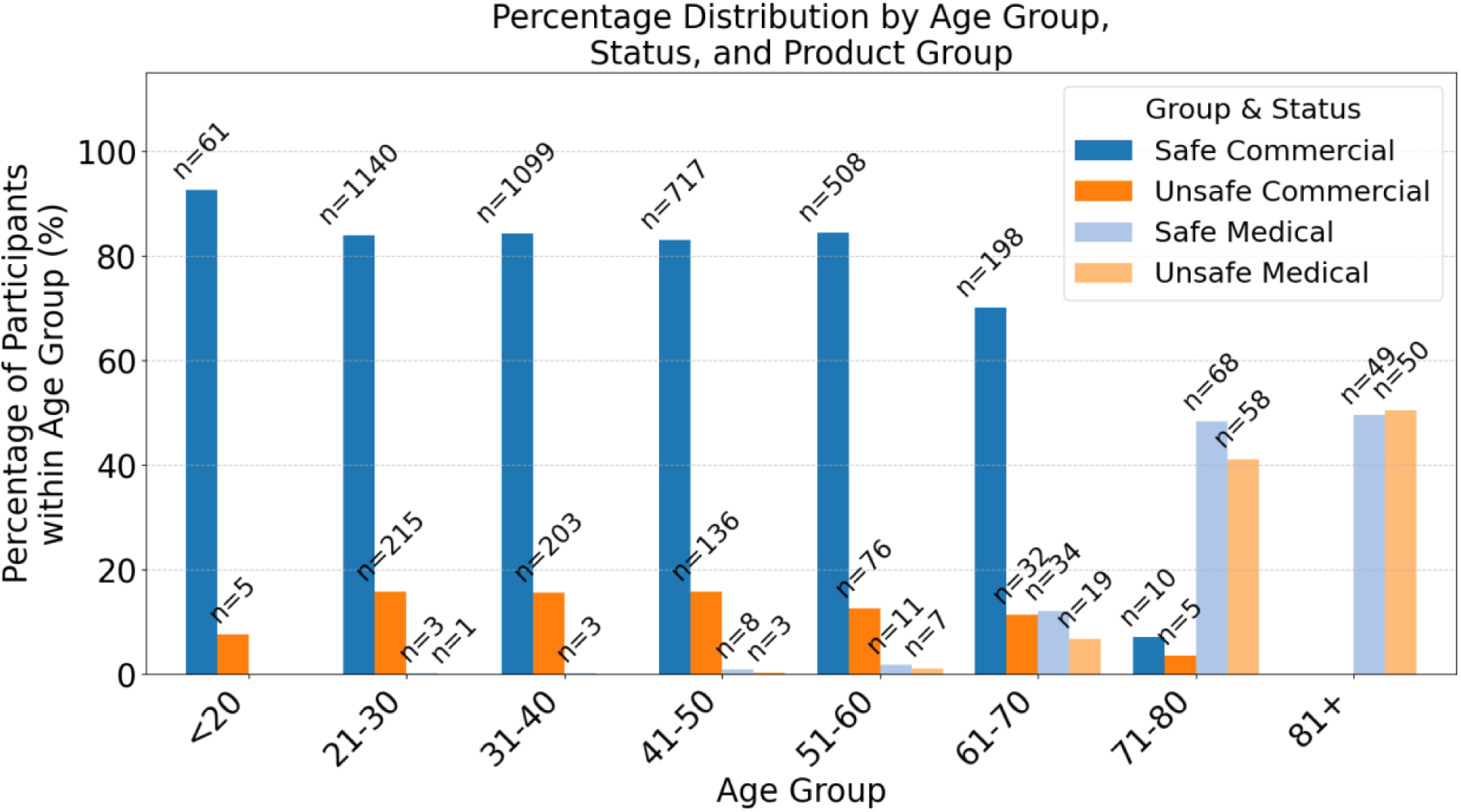
Group percentages by data source and safety group per age in the full dataset (n=4719).

### 2.2 Procedure

Following the general method from Scott et al. (2023), each participant was administered a battery of cognitive tasks related to driving performance (Atkin et al., 2024; Bakhtiari et al., 2020; Scott et al., 2023). The tasks in this study included reaction time (simplified trails), a modified TMT-A, a modified TMT-B, and judgement, memory, and sensorimotor control tasks (Army Individual Test Battery, 1944; Bakhtiari et al., 2020; Reitan, 1958). The previous task battery, Vitals, had four tasks which measured simple reaction time, judgement, working memory, and sensorimotor control (Atkin et al., 2024; Bakhtiari et al., 2020; Scott et al., 2023). Participants in this study completed the latter three Vitals tasks, but they were not analyzed for this study. The Neurapulse battery primarily measures simple reaction time in the Motor task, working memory in the Modified TRT-A and TRT-B tasks, visuomotor tracking in all three tasks, and cognitive flexibility in the Modified TRT-B task (Arbuthnott & Frank, 2000; Crowe, 1998; Kortte et al., 2002; Sánchez-Cubillo et al., 2009). The on-road evaluation was the same as in Scott et al. (2023) and Bakhtiari et al. (2020) for the commercial and medical driving groups respectively, and was designed to test driving ability as judged by a trained instructor.

Most participants were recruited to the study as part of a pre-hire evaluation for prospective employees, and a minority were recruited as a part of referrals to a medical professional where driving risk was suspect (Scott et al., 2023). Individuals that couldn’t complete all three tasks were excluded from the analysis (Table 1). Figures 2 and 3 show the process employed for data collection for both the in-office and on-road assessments, depending on whether collection was from a commercial or medical source, respectively. The data collection process was slightly different between the commercial and medical data sources.

**Figure 2:**
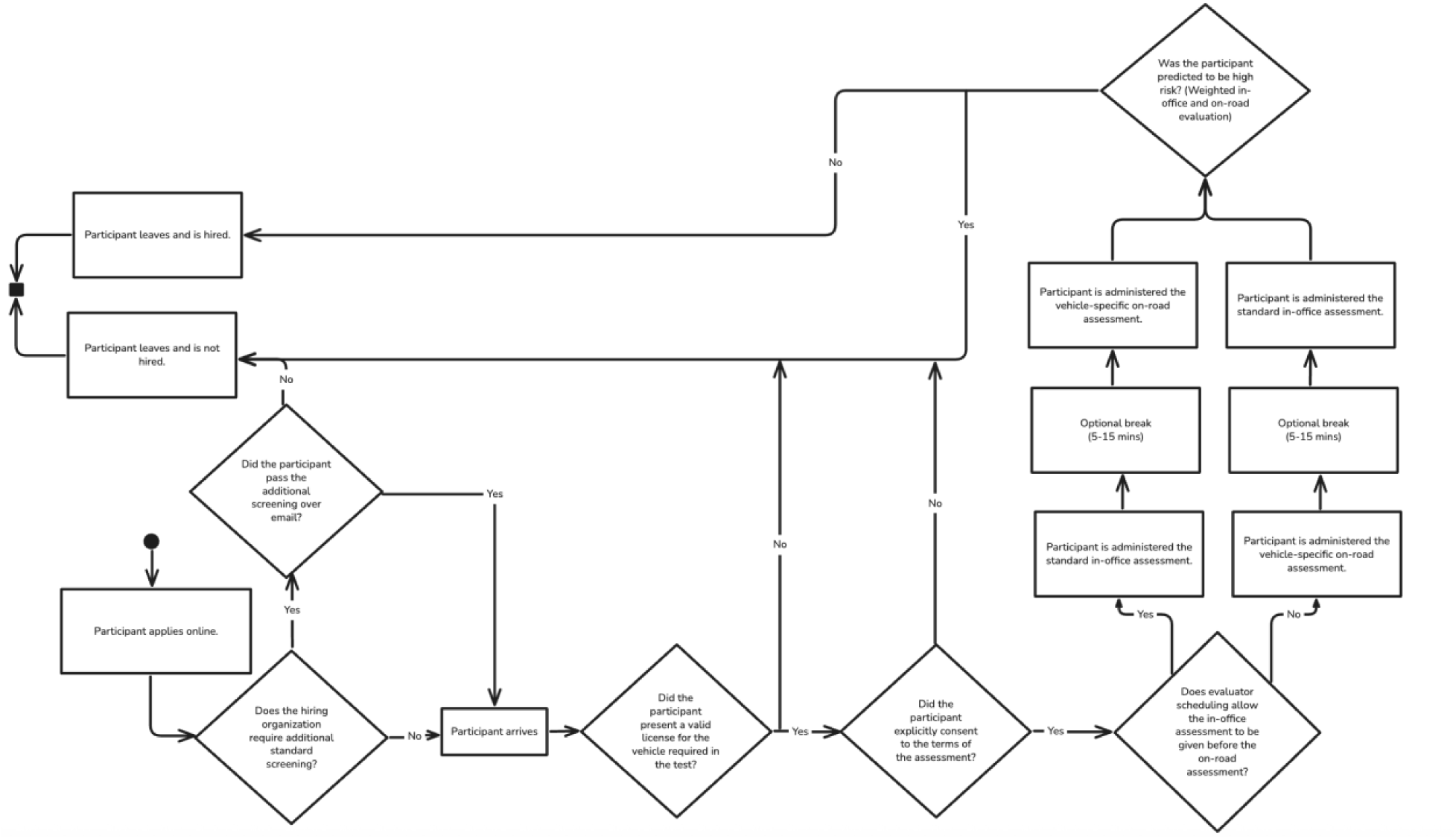
Unified Modelling Language (UML) diagram detailing the participant screening process and decision tree for the on-road and in-office assessment pre-hiring procedure, which was used for commercial data sources. This process is identical to Scott et al. (2023).

**Figure 3:**
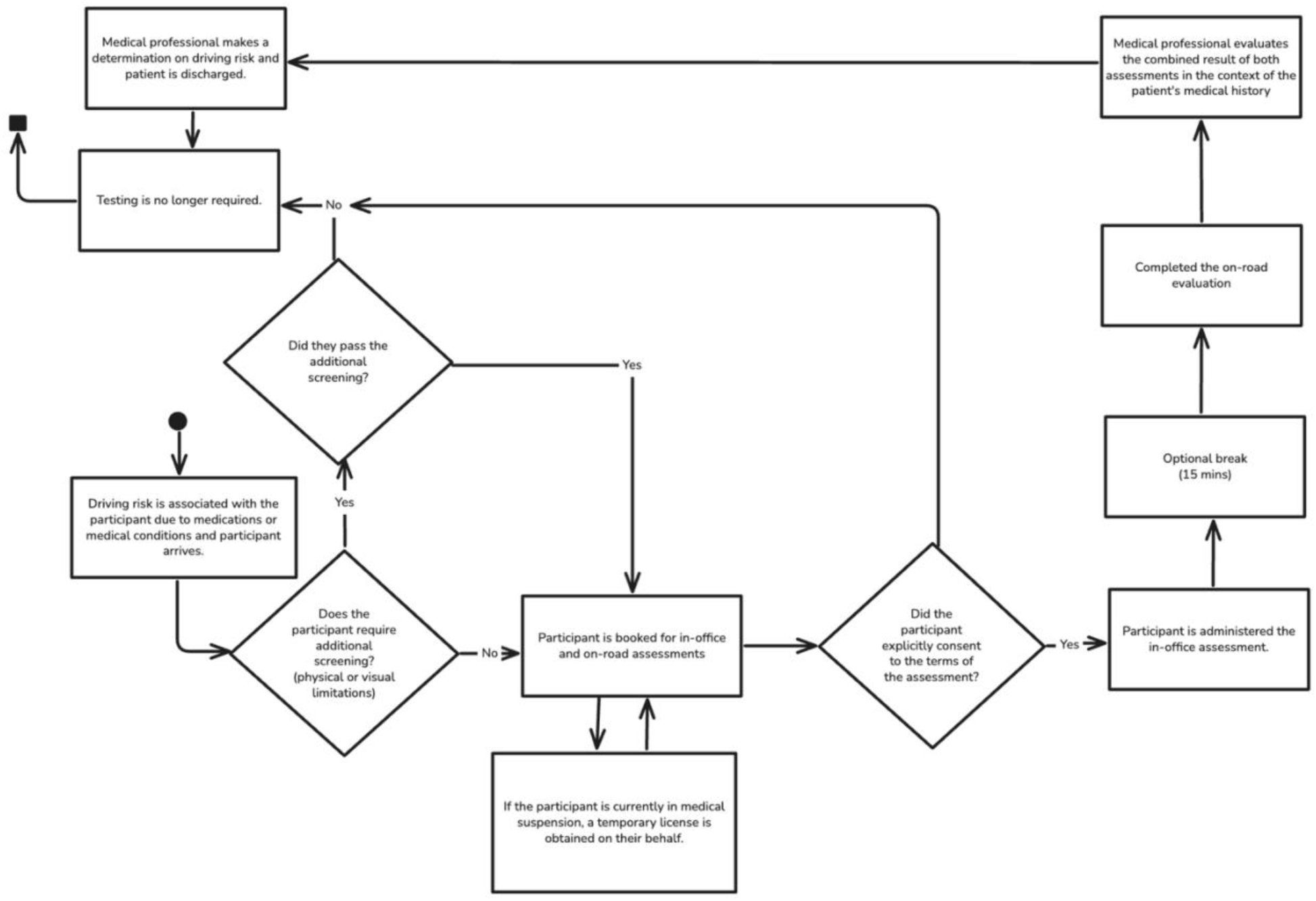
Unified Modelling Language (UML) diagram detailing the participant process and decision tree for the on-road and in-office assessment medical assessment, which was used for medical data sources.

To verify external validity, a logistic regression model with a single, shared cutpoint was used to predict on-road evaluation outcomes in both medical and commercial driving populations. Independent of the data source, an identical task performance was assigned a standardized probability of risk.

### 2.3 Tasks

#### 2.3.1 Cognitive tasks

In our analysis, we included the three Neurapulse tasks that were adapted from the original Trail Making tasks (TMT) and slightly modified by Impirica to run as a continuous ordered series on an electronic device (Army Individual Test Battery, 1944; Bakhtiari et al., 2020; Reitan, 1958; Scott et al., 2023). These 3 tasks included:

1. Reaction Time (Motor): Participants connect an unlabelled set of dots as quickly as possible. After pressing a start button at the bottom of a screen, a dot is shown and the participant must draw a line to that dot with their finger by clicking and dragging a pencil icon. Each time the participant connects to a dot, a new dot is revealed, a previously revealed dot is hidden, but a trail of where the participant has drawn persists for the duration of the stage. This repeats until the maximum number of dots have been shown. There are two stages: a 9-dot configuration and an 18-dot configuration. If the participant lifts their finger before the stage is completed, they are prompted to place their finger back on the pencil icon and continue drawing until the stage is completed. There is no constraint on time.
2. Modified Trails A (TMT-A): Participants connect a sequence of dots as quickly as possible in their correct numerical order. After pressing a start button at the bottom of the screen, a numbered set of dots are shown. The participant must connect the dots in increasing numerical order (1-2-3-4…), starting with dot 1 and ending in the largest number presented by clicking and dragging a pencil icon. There are two stages: a 9-dot configuration and an 18-dot configuration. If the participant lifts their finger before the stage is completed, they are prompted to place their finger back on the pencil icon and continue drawing until the stage is completed. If the participant touches the wrong dot, the pencil icon is moved off the incorrect dot and the participant is prompted to continue. There is no constraint on time.
3. Modified Trails B (TMT-B): Participants connect a sequence of dots as quickly as possible in their correct numerical order. After pressing a start button at the bottom of the screen, two numbered set of dots are shown. One consecutive integer set starts with the number 1, and the other consecutive integer set starts with the number 25. Like TMT-A, each set of numbers is monotonic increasing. participant must connect the dots in increasing numerical order, alternating between each set (1-25-2-26…), starting with dot 1 and ending in the largest number presented by clicking and dragging a pencil icon. There are two stages: a 9-dot configuration and an 18-dot configuration. If the participant lifts their finger before the stage is completed, they are prompted to place their finger back on the pencil icon and continue drawing until the stage is completed. If the participant touches the wrong dot, the pencil icon is moved off the incorrect dot and the participant is prompted to continue. There is no constraint on time.

Shorter TMT tasks are capable of measuring the same aspects of cognition as the longer 25-target paper and pencil tests (Dahmen, 2017). In our Modified TMT, each of these tasks uses a 9-dot and 18-dot configuration to adapt better for smaller screen sizes. In this configuration, the task battery is designed to take under 5 minutes.

#### 2.3.2 Dot Configuration of the Trails Tasks

The originally validated TMT task from Reitan (1958) suffers from a lack of effectiveness in longitudinal cognitive assessment due to the influence of practice effects on sensitivity (Zeng et al., 2017). To increase the effectiveness of test-retest reliability, the dot coordinates in our version of the task are algorithmically generated using a modified version of the algorithm created by Zeng et al. (2017) in a manner that preserves the spatial characteristics of Reitan’s configuration of TMT and maintains equivalent predictive power under the proposed regime. Our version of the task uses the divide-and-combine (DAC) approach from Zeng et al. (2017) but required an extra phase of applying padding and jitter on final dot coordinates to avoid overlaps once rescaled across the range of aspect ratios found on digital devices. Using the DAC approach alone, we found that there was a high number of generations which had either no solution due to path intersections or had very asymmetric density. Pathing intersection issues included cases where either A) at least one dot intersected with at least one path-correct line segment in at least one supported aspect ratio; or B) at least one dot or at least one line intersected with the position of the start button. To satisfy support for a range of aspect, a small amount of jitter was applied repeatedly to dot coordinates until pathing criteria were satisfied. This jitter phase was bounded between 1 and 20000 steps, so if the number of steps increased beyond 20000, the dot configuration candidate was rejected and regenerated. Under this modification, we generated a set of 100 algorithmically generated solutions that matched the spatial characteristics demonstrated by Zeng et al. (2017) but found the coordinate solutions to be more robust across a range of screen sizes. When a participant runs a Motor or TMT Task, one of the 100 solutions are randomly drawn and presented.

#### 2.3.3 Extension to Zeng et al. (2017) Method

Formally, the steps to apply jitter are generally as follows:

1. The coordinates are mapped from their originally generated domain under the DAC regime [1000,2000] to [-0.5,0.5] for both x and y axes.
2. The core jitter update rule modifies each target position by adding a random displacement vector where *p_i* represents the 2D position vector of target *i* at iteration *t*, and *ε_i* is the corresponding jitter vector:

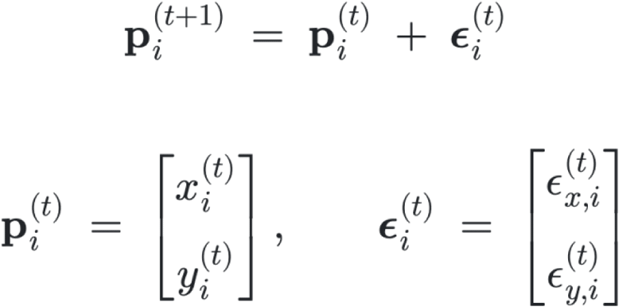
3. The jitter components are sampled independently from Gaussian distributions with different standard deviations for horizontal and vertical displacement. In our case, the vertical jitter has a larger standard deviation (0.07 vs 0.02) to provide more movement flexibility in the y-axis (avoiding intersection with the start button at the bottom), but these values can be set to 0.02 for both *x* and *y* for general cases. The following values assume portrait orientation and an initial 2:1 aspect ratio:

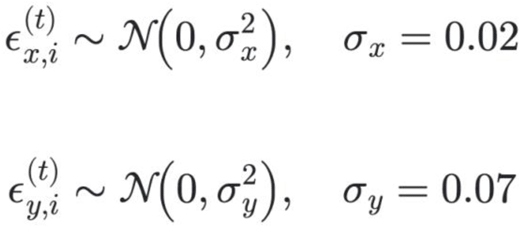
4. After applying the jitter, positions are constrained to remain within the valid coordinate space:

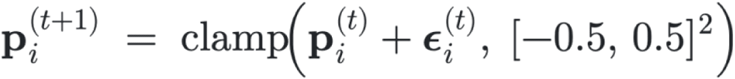
5. The algorithm terminates when the collision constraint function reaches zero, indicating no remaining conflicts in either the base or transformed scale:

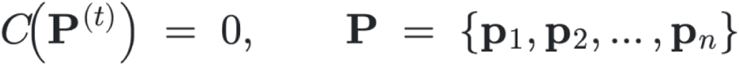

where *C* is the collision detection function that counts the total number of constraint violations across all target positions *P in both scales*. In our case, this function checks the count of any intersections between line segments, dots, and bounding boxes.

**Figure 4:**
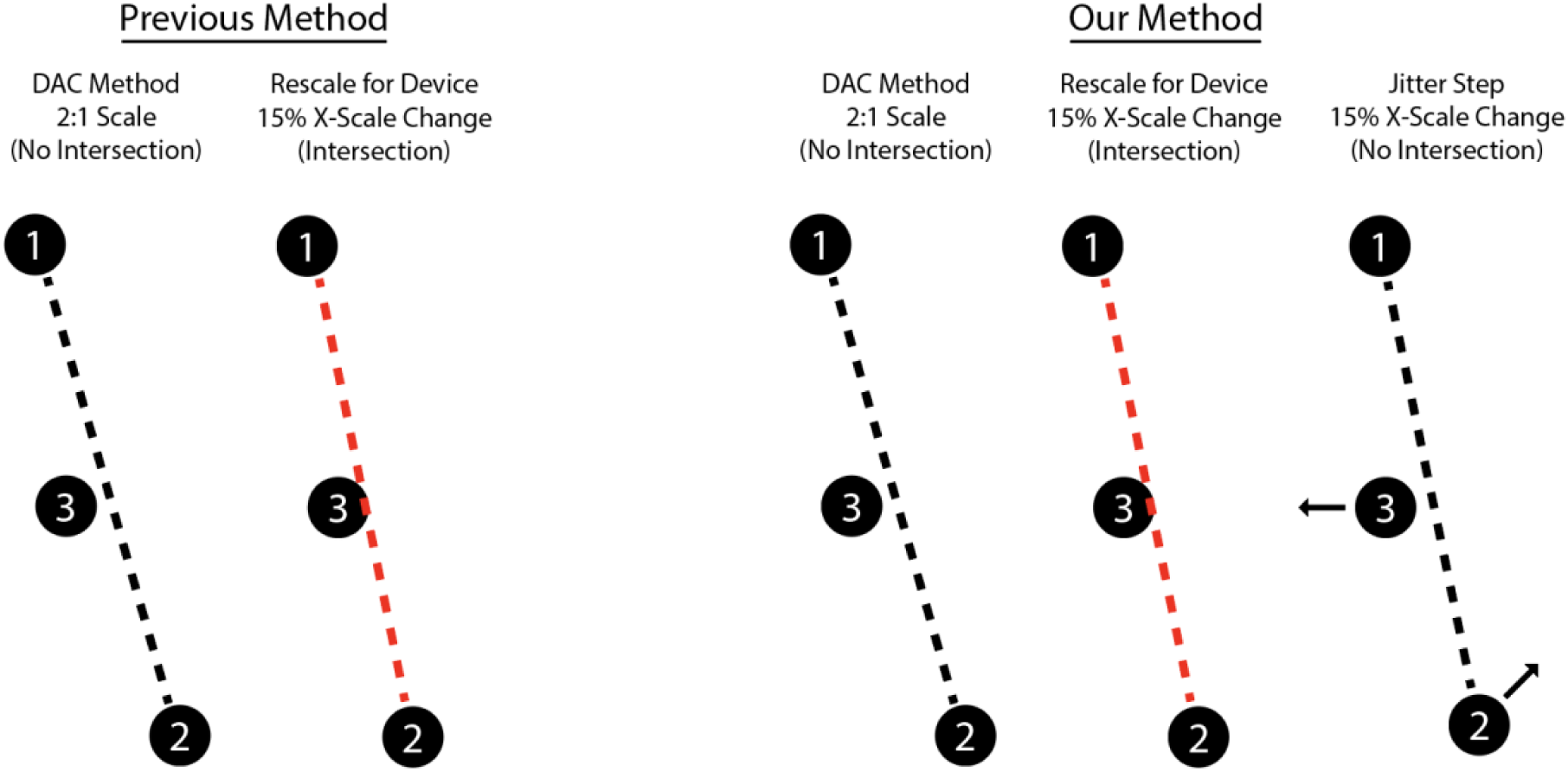
Illustration of the pathing issues that can appear during TMT configuration using the Zeng et al. (2017) method (left), and our improved method for stabilizing solutions across a range of aspect ratios (right).

**Figure 5:**
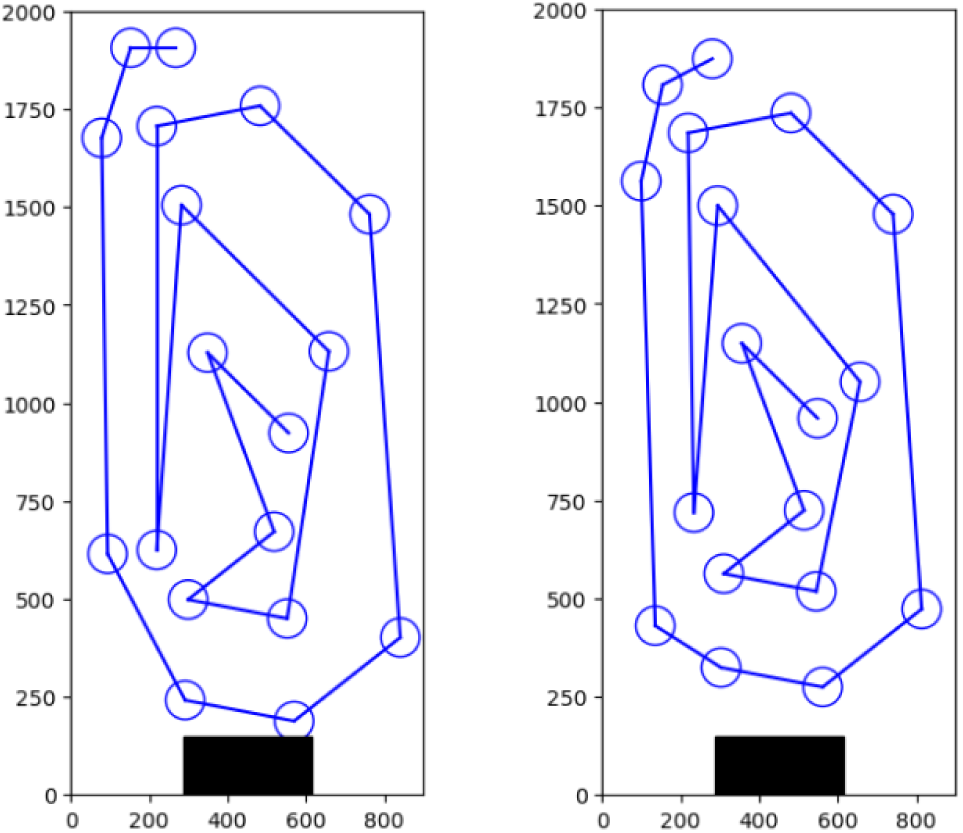
Example of the intersection issues that can appear during TMT configuration using the Zeng et al. (2017) method (left), and after our improved method (right) which includes a jitter step to fix intersections that violate the spatial configuration of the original TMT task. The bounding box for the start button is seen in black.

#### 2.3.4 Dual Number Sequence of the Modified TMT-B

The TMT task has been widely validated in literature but is culturally loaded due to its reliance on the Latin alphabet (Bezdicek et al., 2016; Zhao et al., 2013). To address this, many researchers have presented variants to the standard TMT-B sequence (1-A-2-B-) which include the Shape Trail Test (STT-B), the Colors Trails Test (CTT-B), and the Black and White TMT (TMT-B&W) (D’Elia et al., 1996; Kim et al., 2014; Zhao et al., 2013). The TMT variants have limitations including the CTT-B’s pink-yellow color scheme which may bias performance in cases of Tritanopia (National Eye Institute, 2023) and is significantly more cognitively loading than TMT-B (Dugbartey, 2000), the TMT-B&W which is significantly less cognitively loading than the original TMT-B (Friberg, 2017), and the SST-B which is more cognitively loading than CTT-B (Zhao et al., 2013). For culture-fairness, we use two sets of numerical sequences (1-25-2-26-) instead of TMT-B (1-A-2-B-) in our Modified TMT-B task, which tests the cognitive flexibility required for switching between sequences while avoiding effects from differences in cultural salience due to letters or shapes.

**Figure 6:**
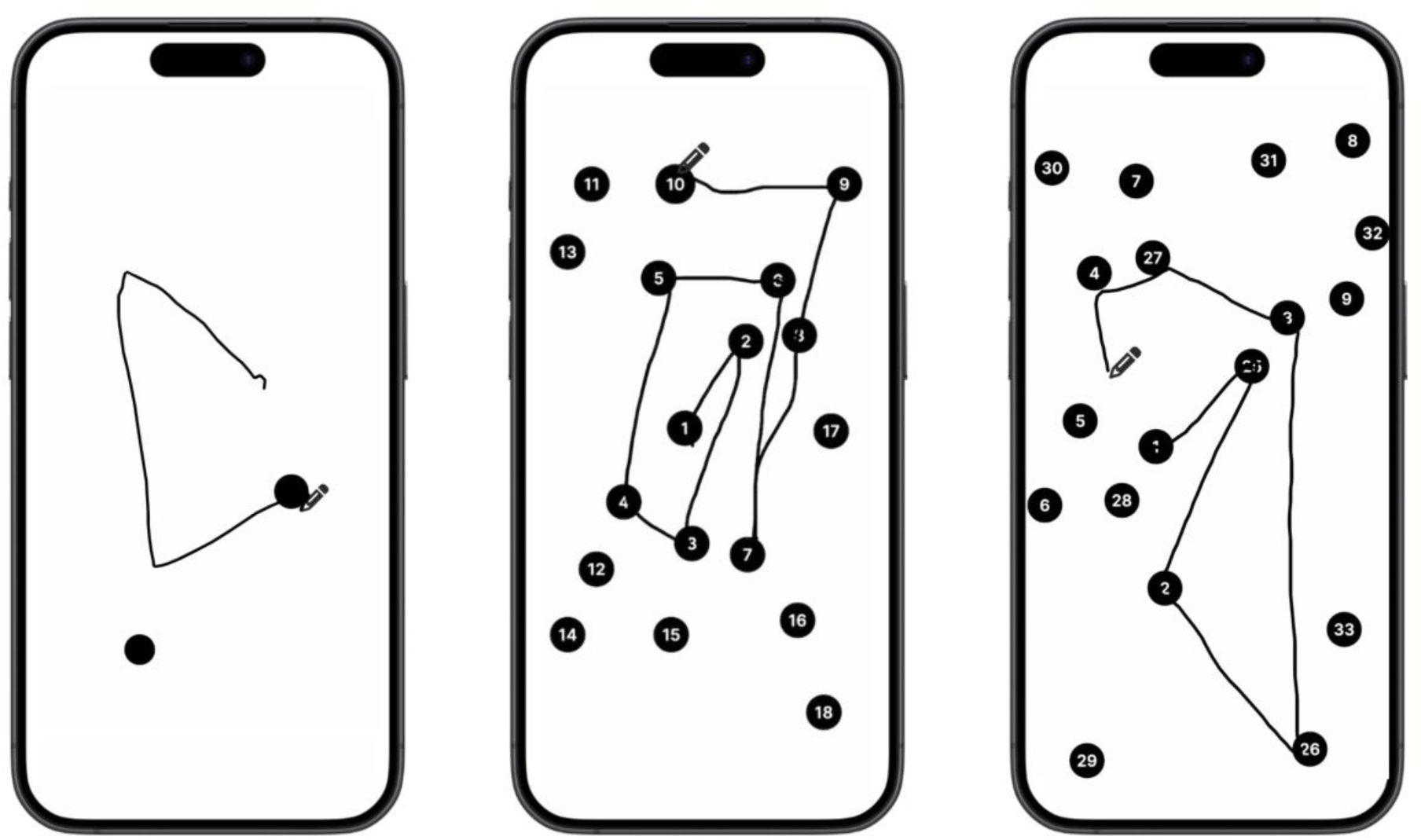
Example of a partially completed 18-target Motor stage, 18-target Modified TRT-A, and 18-target Modified TRT-B stage on a 2.13:1 portrait aspect ratio digital device (left to right).

#### 2.3.5 On-Road Evaluation

In our study, participants from commercial data sources used the Commercial On-Road Evaluation (CORE) for on-road evaluation, and participants from medical data sources used the DriveABLE On-Road Evaluation (DORE) for on-road evaluation. The DORE was originally developed by Dobbs et al. (1998) and was optimized to distinguish between the types of driving errors made by cognitively intact drivers versus those with mild-cognitive impairment (MCI) or Alzheimer’s Disease (AD). Unlike driver’s license evaluations, the DORE tests the cognitive skills needed for safe driving (Berndt et al., 2007; Dobbs et al., 1998; Kowalski and Tuokko, 2007). For the on-road score produced by these evaluations, the penalty is weighted towards driving errors that are made by drivers in the cognitively at-risk groups. The DORE score is used by medical professionals as an objective measure of cognitive risk as it relates to driving in combination with the driver’s medical record. The CORE is adapted from the DORE, calibrated to the same errors but adapted for commercial vehicles. This is due to physical differences in vehicle size and controls, including brakes, maneuverability, blind spots, and necessary visual scanning. The total difference in errors between vehicle types is 4% (three differences across 70 error categories; Scott et al., 2023). The on-road evaluation is administered by a driving evaluator who is certified by a standardized training program at Impirica. Using the methodology of Scott et al. (2023), drivers that score as high risk in either assessment are labeled as “Unsafe”, otherwise they are labeled as “Safe” (Table 2).

### 2.4 Neurapulse Dependent Measures

For both the 9-dot and 18-dot levels of the three Neurapulse tasks (Motor, TMT-A and TMT-B), we measured the sum of the log-transformed reaction time (Target Log RT) and the sum of errors that were committed (Target Errors). Target Log RT did not include the time taken to reach the first dot. Target Errors were defined as any instance when the digital pencil was dragged onto the incorrect next dot.

**Table 3:**
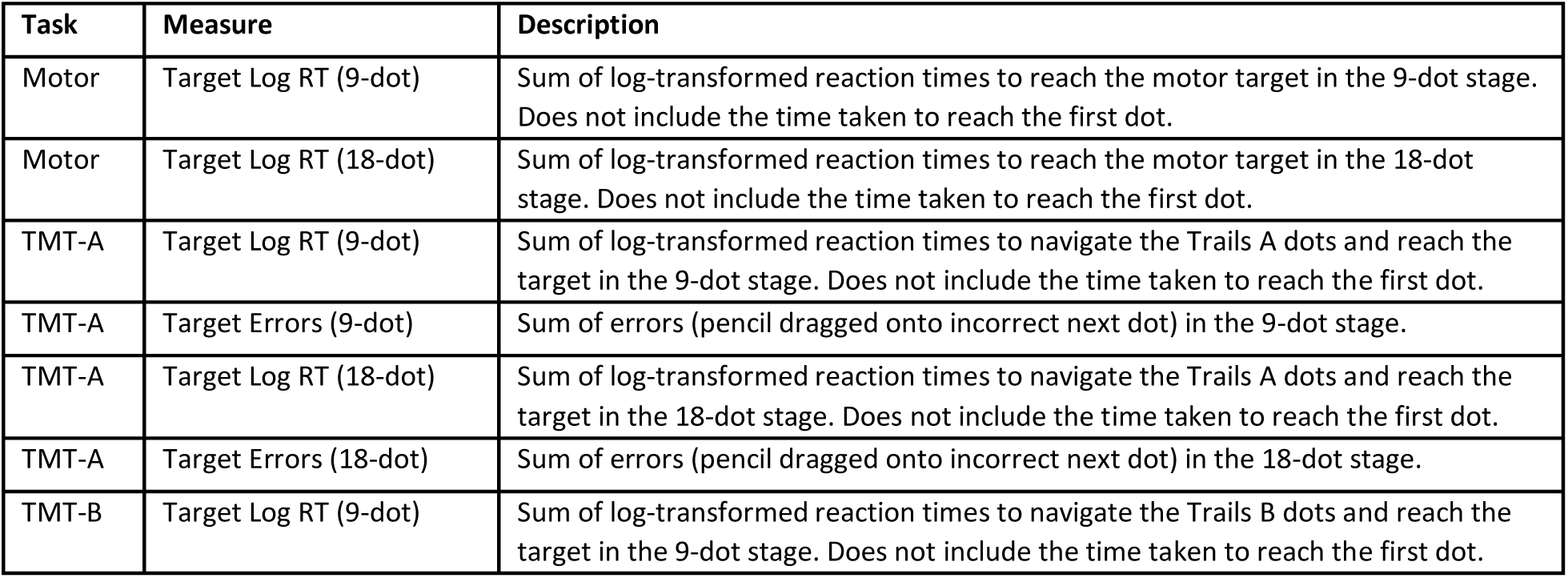
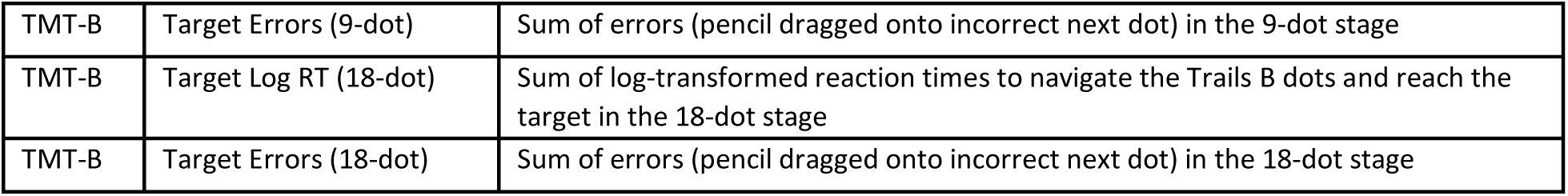
Tasks and dependent measures in the Neurapulse battery.

### 2.5 Statistical Analysis

Participants were divided into two equally sized groups for model training (n=2360) and testing (n=2359). Within each split, the number of samples obtained from each data source was split equally (Figure 7) The training group contained 157 participants from medical and 2203 participants from commercial data sources. The testing group contained 157 participants from medical and 2202 participants from commercial data sources. For variable selection criteria, we included candidates which showed significant post hoc differences between safe and unsafe driving outcomes across all data sources (total time) and were not considered to be a practice trial for the participant (18 target). This resulted in a total of 3 variables for our model candidates: the sum of log-transformed time in the 18-target Motor task, the sum of log-transformed time in the 18-target Modified TMT-A task, and the sum of log-transformed time in the 18-target Modified TMT-B task. Instead of using the sum of time across each of the tasks, we use the sum of log-transformed reaction times to normalize their distributions (Whelan, 2008).

**Figure 7:**
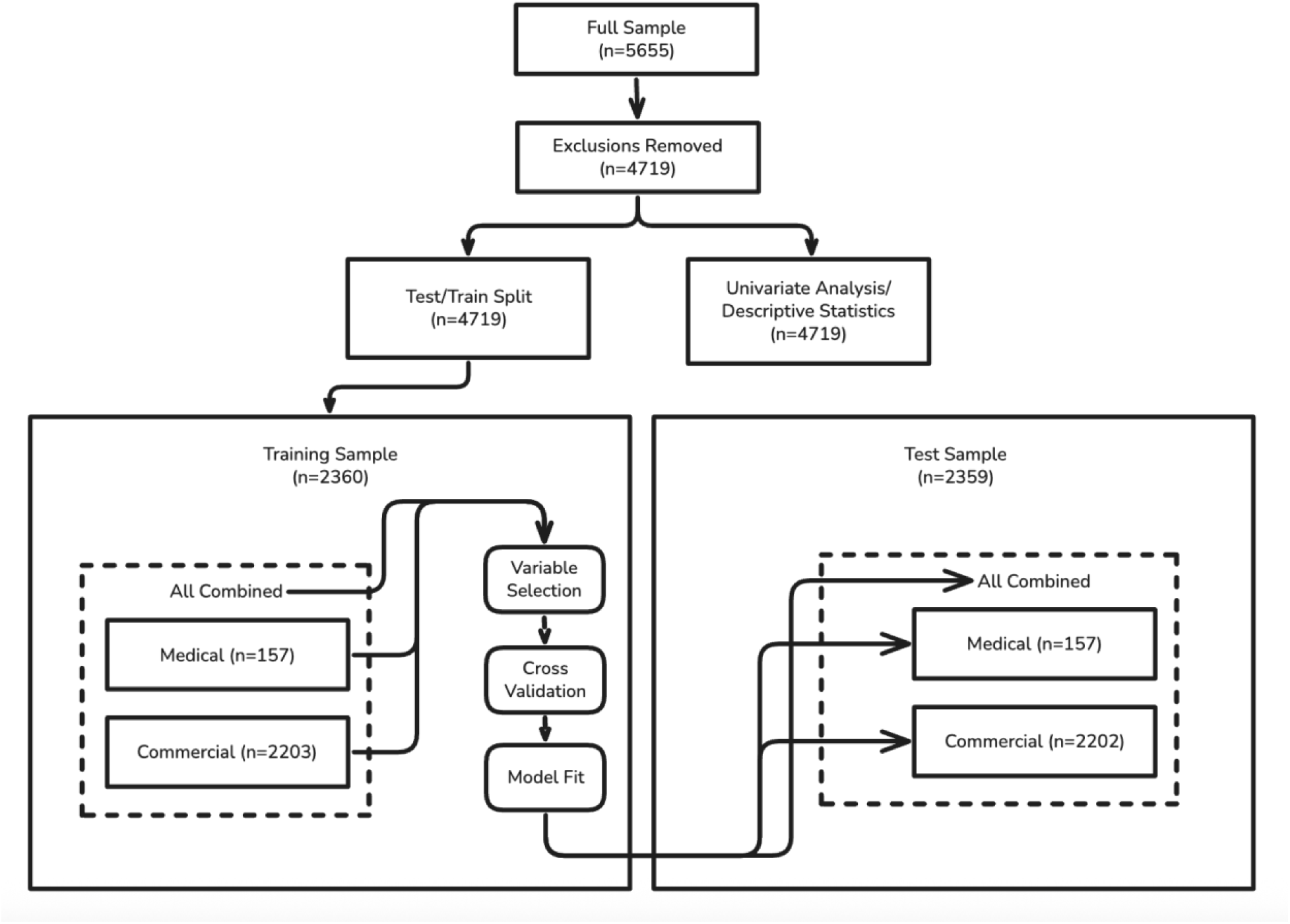
Training and test data split participant counts.

### 2.6 Stratification

We fit an additional set of models using stratified training data using a 2×2 blocking matrix, to see if model performance in the held-out test set improved or worsened. The matrix was configured as driving outcome (safe and unsafe) and data source (medical and commercial), and then randomly oversampled (Batista et al., 2004; He & Garcia, 2009). Large age ranges were missing within some of the subgroups and therefore were not suitable to be included as a stratification group. For example, there was a lack of younger participants available within the medical data source (See Figure 1). The stratification procedure included counting the participants in each subgroup, identifying the largest group count (safe, commercial, n=1867), and then randomly sampling from the original subgroup with replacement until the group was in equivalent in size to the largest group (Table 4).

**Table 4:** Training dataset sizes before and after stratification.

| Safety Status | Group | Count Before | Count After |
| --- | --- | --- | --- |
| Safe (0) | Commercial | 1867 | 1867 |
| Safe (0) | Medical | 82 | 1867 |
| Unsafe (1) | Commercial | 336 | 1867 |
| Unsafe (1) | Medical | 75 | 1867 |
| Total | Total | 2360 | 7468 |

### 2.7 Model Fitting Method

Using the training data split, we analyzed a logistic regression model under the L2 normalization regime across all data sources using the scikit-learn and statsmodels python packages in the same manner as Scott et al. (2023) and Atkin et al. (2024), and included all 3 variables that met the selection criteria (Pedregosa et al., 2011, Scott et al., 2023, Seabold & Perktold, 2010). The model fit is reported for all combined data in addition to each data source group to support external validity.

### 2.8 Accuracy Methods

Classifying the paired in-office and on-road test outcomes in the manner of Scott et al. (2023), if the participant was classified as “safe” by the model and passed the on-road road course then they were labelled as a true negative (TN), if the participant was classified as “safe” by the model and failed the on-road road course then they were labelled as a false negative (FN), if the participant was classified as “unsafe” by the model and passed the on-road road course then they were labelled as a false positive (FP), and if the participant was classified as “unsafe” by the model and failed the on-road road course then they were labelled as a true positive (TP) (Scott et al., 2023). The positive predictive value (PPV) measures the probability that a participant failed the on-road test if the model classified them as “unsafe”, the negative predictive value (NPV) measures the probability that a participant passed the on-road test if the model classified them as “safe”, and accuracy (ACC) measures the proportion of cases that the model correctly predicted the on-road test outcome.

**Figure 8:**
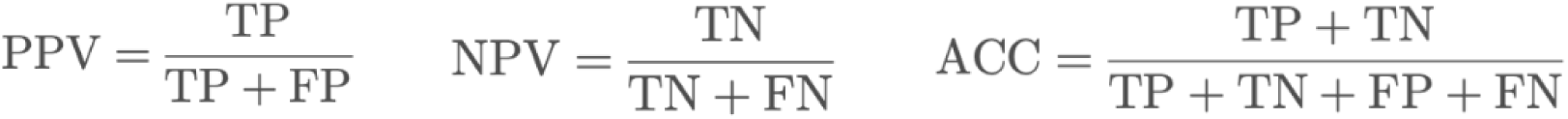
Formulas for positive predictive value, negative predictive value, and accuracy, respectively.

**Figure 9:**
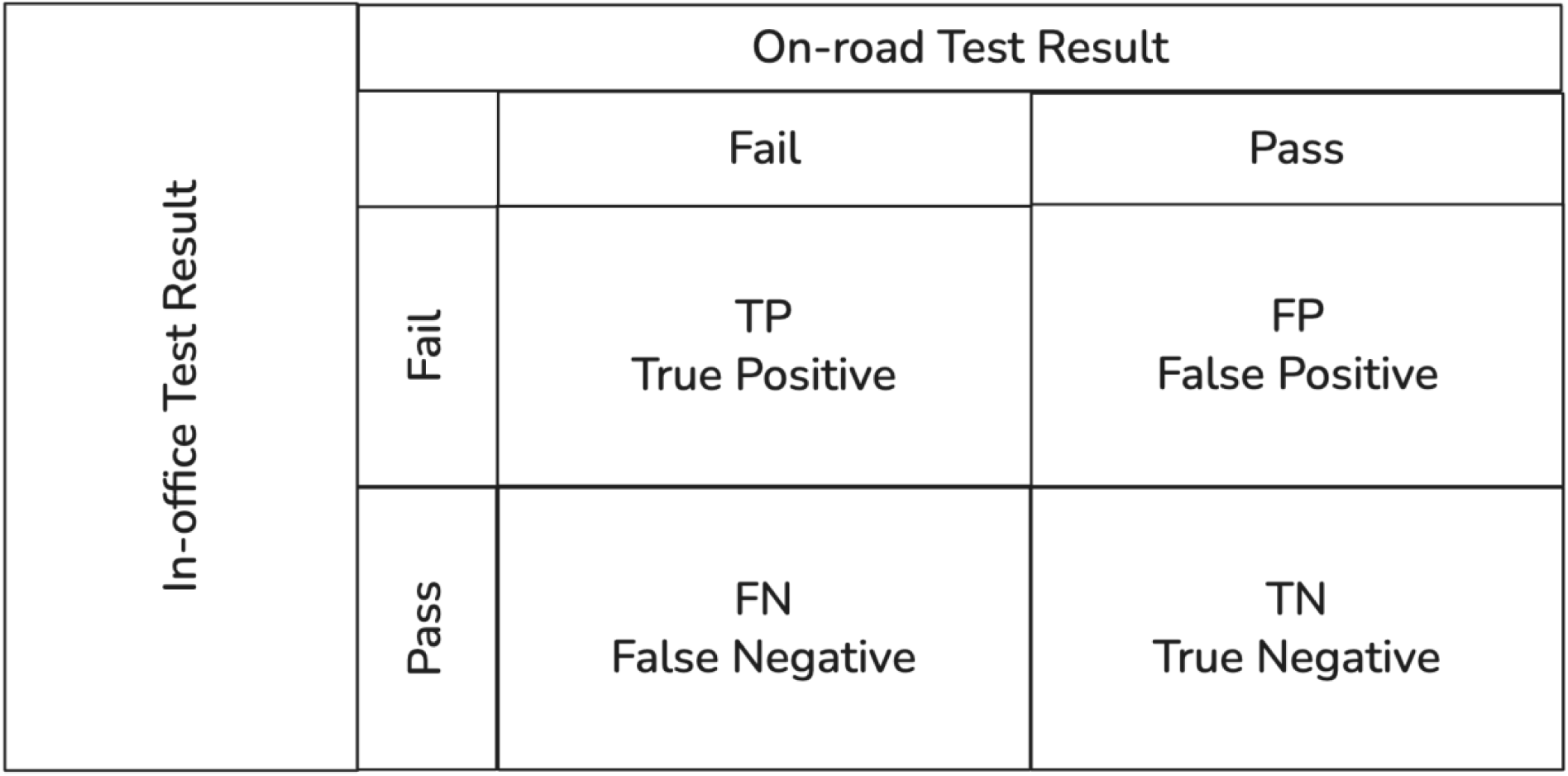
Confusion matrix definition for in-office test and on-road test results.

## 3. Results

### 3.1 Descriptive Statistics

Table 5 describes the differences in performance between all participants that were classified safe or unsafe in the standard on-road assessment, including the mean, median, and standard deviation for each group. Following the methodology of Scott et al. (2023), if a participant was found to be high risk under the on-road assessment scoring regime, they were classified as unsafe, otherwise they were classified as safe. (Scott et al., 2023).

**Table 5:** Mean and median performance on Neurapulse tasks by safety group in the medical dataset (n=314).

| Metric | Mean Safe | Mean Unsafe | Median Safe | Median Unsafe | Std Safe | Std Unsafe | Min Safe | Min Unsafe | Max Safe | Max Unsafe |
| --- | --- | --- | --- | --- | --- | --- | --- | --- | --- | --- |
| Motor 9 Target Log RT | 3.464 | 4.923 | 2.776 | 4.252 | 2.983 | 3.529 | -1.795 | -1.456 | 12.123 | 18.898 |
| Motor 18 Target Log RT | 0.577 | 3.267 | 0.214 | 2.393 | 3.853 | 5.450 | -6.714 | -5.530 | 11.239 | 26.036 |
| Trails A 9 Target Log RT | 3.431 | 5.024 | 3.082 | 4.099 | 2.952 | 3.889 | -3.604 | -1.512 | 17.208 | 23.257 |
| Trails A 18 Target Log RT | 5.769 | 9.279 | 4.851 | 7.780 | 7.059 | 7.884 | -12.318 | -7.779 | 34.447 | 35.033 |
| Trails B 9 Target Log RT | 9.345 | 15.342 | 7.403 | 10.889 | 7.458 | 21.309 | -0.858 | -1.426 | 54.132 | 209.835 |
| Trails B 18 Target Log RT | 22.165 | 29.656 | 18.526 | 23.153 | 15.165 | 22.161 | -0.087 | -3.520 | 84.133 | 118.308 |
| Trails A 9<br>Target Errors | 0.415 | 0.659 | 0.000 | 0.000 | 0.928 | 1.287 | 0.000 | 0.000 | 6.000 | 8.000 |
| Trails A 18<br>Target Errors | 0.909 | 1.312 | 0.000 | 1.000 | 2.238 | 2.499 | 0.000 | 0.000 | 19.000 | 20.000 |
| Trails B 9<br>Target Errors | 1.443 | 3.478 | 0.000 | 1.000 | 2.783 | 9.025 | 0.000 | 0.000 | 19.000 | 82.000 |
| Trails B 18<br>Target Errors | 3.250 | 5.167 | 1.000 | 2.000 | 5.954 | 10.388 | 0.000 | 0.000 | 50.000 | 67.000 |

**Table 6:** Mean and median performance on Neurapulse tasks by safety group in the commercial dataset (n=4405).

| Metric | Mean Safe | Mean Unsafe | Median Safe | Median Unsafe | Std Safe | Std Unsafe | Min Safe | Min Unsafe | Max Safe | Max Unsafe |
| --- | --- | --- | --- | --- | --- | --- | --- | --- | --- | --- |
| Motor 9<br>Target Log RT | 0.155 | 0.560 | -0.094 | 0.242 | 1.922 | 2.184 | -4.458 | -4.822 | 17.085 | 14.732 |
| Motor 18<br>Target Log RT | -4.486 | -3.877 | -5.000 | -4.530 | 3.424 | 3.966 | -14.079 | -12.057 | 19.473 | 20.930 |
| Trails A 9<br>Target Log RT | 0.168 | 0.803 | 0.131 | 0.654 | 2.236 | 2.446 | -7.188 | -5.174 | 14.923 | 16.688 |
| Trails A 18<br>Target Log RT | -1.655 | -0.049 | -1.941 | -0.810 | 4.982 | 7.119 | -21.596 | -16.839 | 59.971 | 112.805 |
| Trails B 9<br>Target Log RT | 3.083 | 4.492 | 2.434 | 3.170 | 4.530 | 6.416 | -5.921 | -7.614 | 85.338 | 59.635 |
| Trails B 18<br>Target Log RT | 6.961 | 9.214 | 5.704 | 7.711 | 7.855 | 9.526 | -13.543 | -12.729 | 84.613 | 82.782 |
| Trails A 9<br>Target Errors | 0.137 | 0.174 | 0.000 | 0.000 | 0.485 | 0.626 | 0.000 | 0.000 | 12.000 | 10.000 |
| Trails A 18<br>Target Errors | 0.448 | 0.688 | 0.000 | 0.000 | 1.571 | 4.323 | 0.000 | 0.000 | 40.000 | 97.000 |
| Trails B 9<br>Target Errors | 0.593 | 1.171 | 0.000 | 0.000 | 2.545 | 4.152 | 0.000 | 0.000 | 63.000 | 59.000 |
| Trails B 18<br>Target Errors | 1.298 | 2.225 | 0.000 | 0.000 | 4.820 | 7.209 | 0.000 | 0.000 | 96.000 | 89.000 |

### 3.2 Group Comparisons

Participant performance on the Motor, Modified TMT-A, and Modified TMT-B tasks were compared on the levels of safety status (pass, fail) and data source (medical, commercial). In both medical and commercial data sources, participants that were classified as unsafe on the on-road course had a significantly slower completion time on all three in-office tasks compared to drivers that were classified as safe. The number of errors made in the in-office assessment in the Modified TMT-B task, instances where the participant connected a dot outside of the correct sequence, was statistically significant between drivers classified as safe and unsafe for both 9-target and 18-target configurations, but only in the commercial data source. For participants in the medical data source, there was only a significant difference in error count for the 9-target configuration.

On the Motor 9-target task, participants from the medical and commercial data sources had slower reaction times if they were classified as unsafe in the on-road assessment compared to if they were classified as safe (Figure 10). Likewise, on the Motor 18-target task, participants from the medical and commercial data sources had slower reaction times if they were classified as unsafe in the on-road assessment compared to if they were classified as safe.

**Figure 10:**
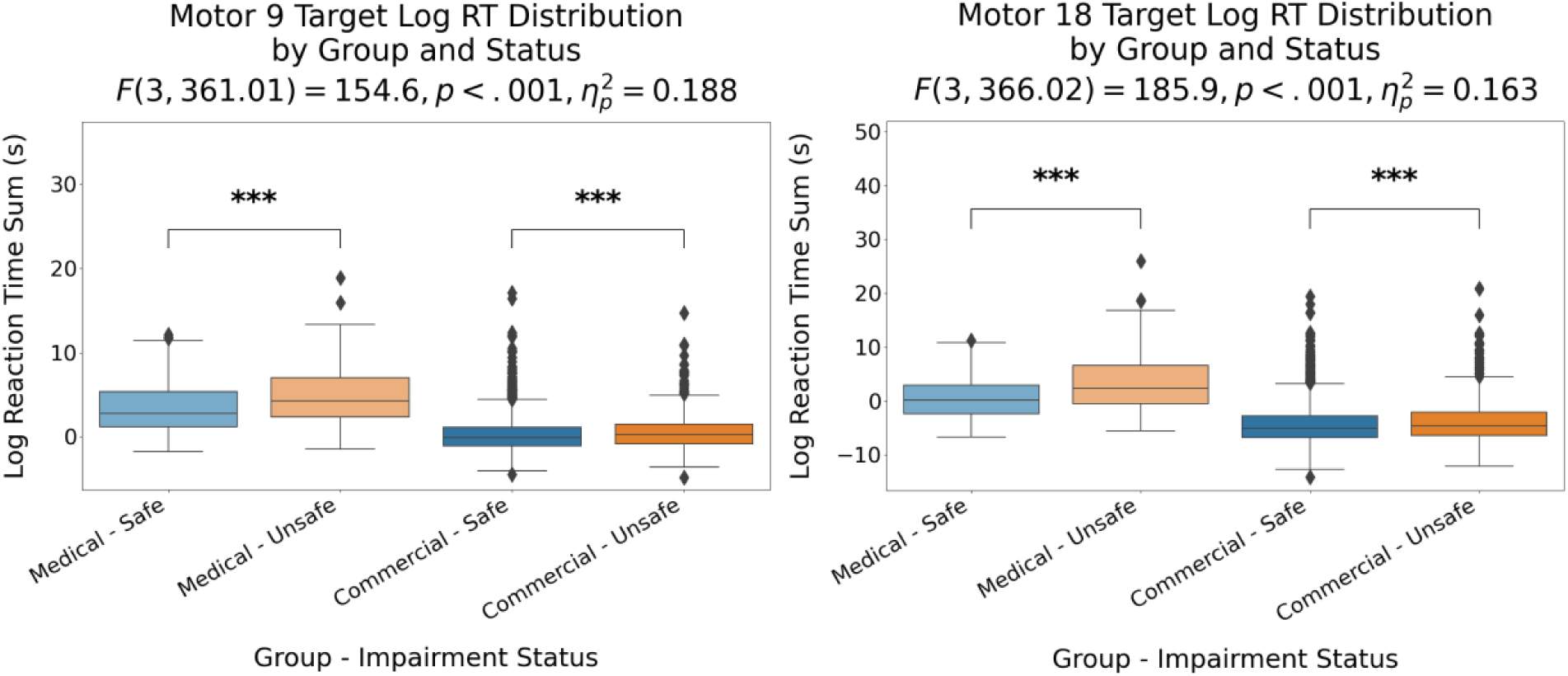
Group means and differences comparing the sum of log reaction times (seconds) on the Motor task depending on data source (medical vs commercial), on-road test outcome (safe vs unsafe), and stage of the Motor task (9 target vs 18 target). Vehicle drivers who failed the road test were slower on the Motor task than drivers who passed the road test.

On the Modified TMT-A 9-target task, participants from the medical and commercial data sources had slower reaction times if they were classified as unsafe in the on-road assessment compared to if they were classified as safe (Figure 11). Likewise, on the Modified TMT-A 18-target task, participants from the medical and commercial data sources had slower reaction times if they were classified as unsafe in the on-road assessment compared to if they were classified as safe.

**Figure 11:**
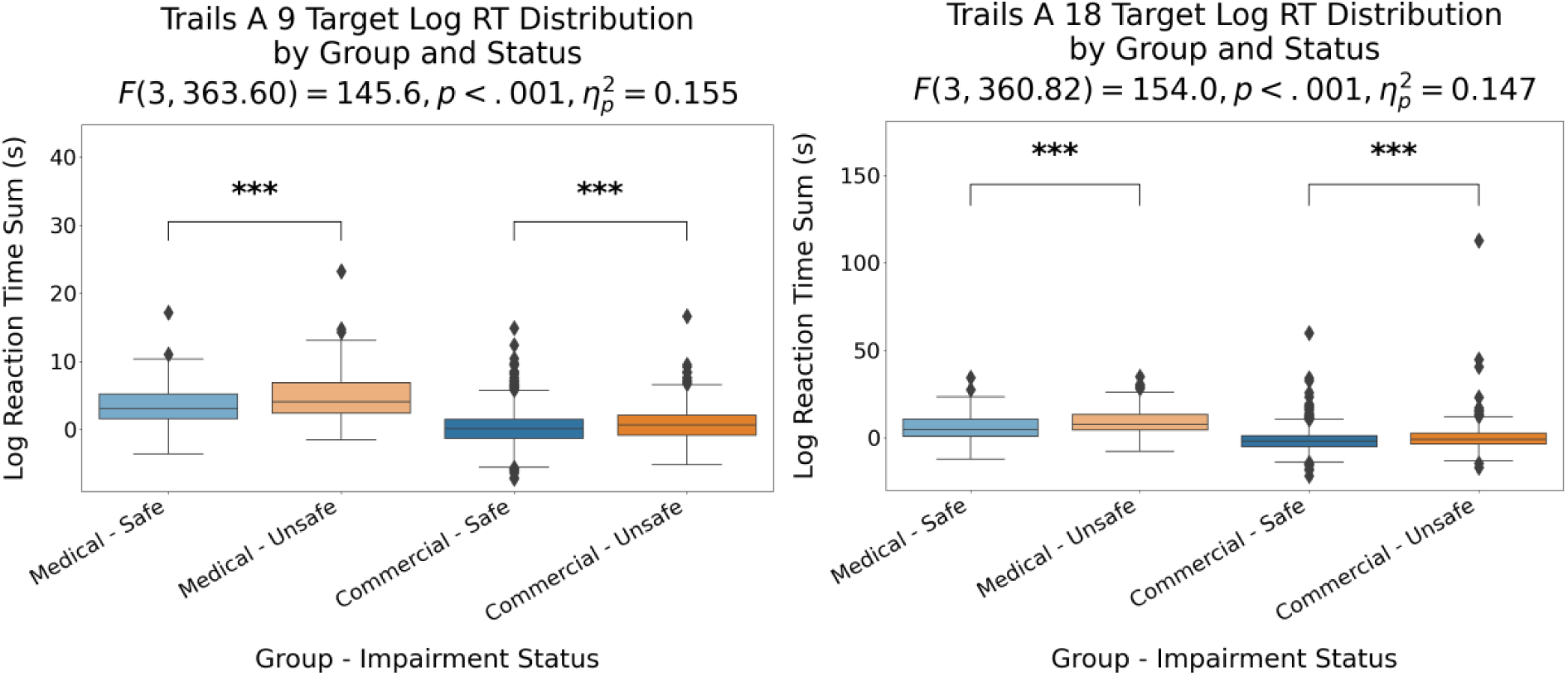
Group means and differences comparing the sum of log reaction times (seconds) on the Trails A task depending on data source (medical vs commercial), on-road test outcome (safe vs unsafe), and stage of the Trails A task (9 target vs 18 target). Vehicle drivers who failed the road test were slower on the Trails A task than drivers who passed the road test.

On the Modified TMT-B 9-target task, participants from the medical and commercial data sources had slower reaction times if they were classified as unsafe in the on-road assessment compared to if they were classified as safe (Figure 12). Likewise, on the Modified TMT-B 18-target task, participants from the medical and commercial data sources had slower reaction times if they were classified as unsafe in the on-road assessment compared to if they were classified as safe.

**Figure 12:**
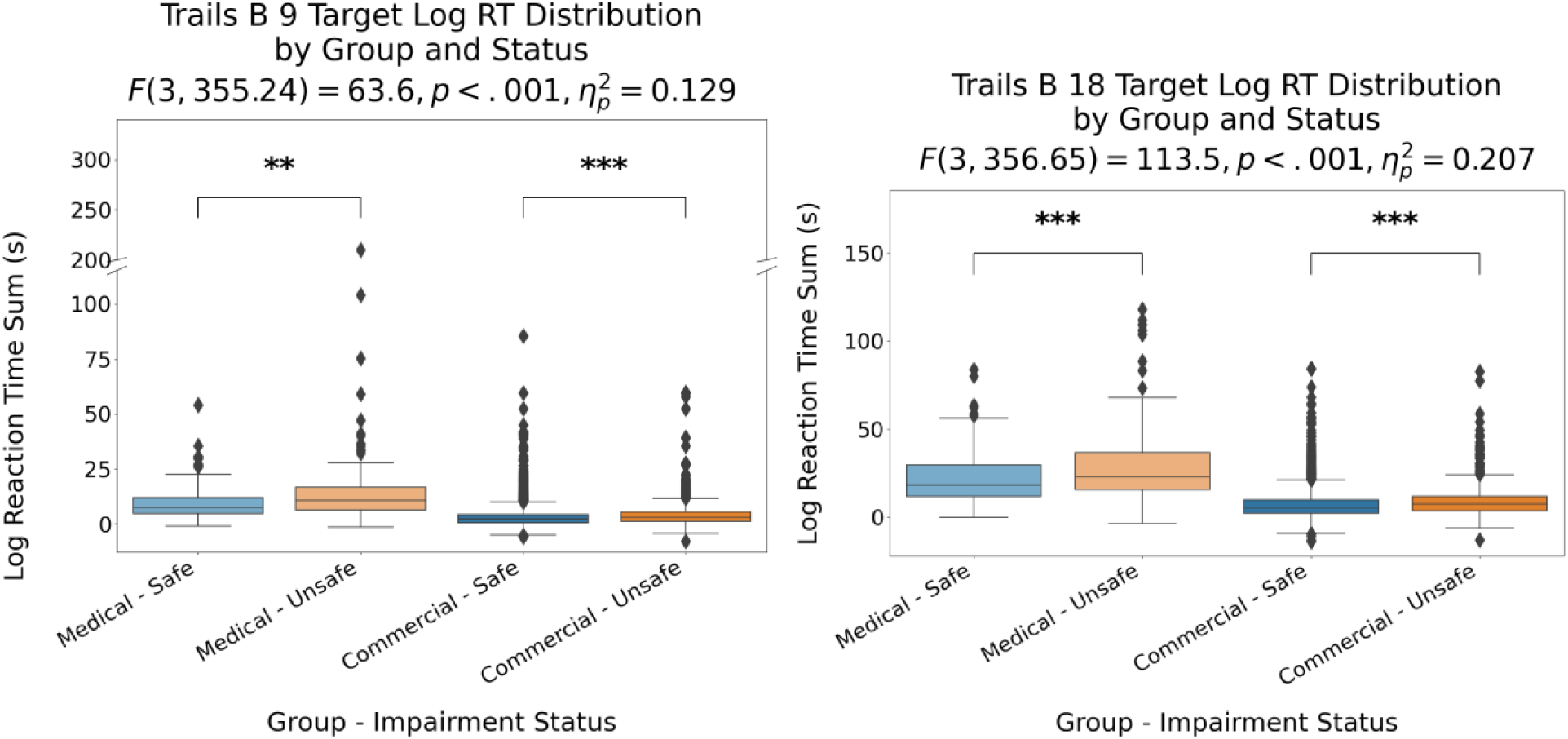
Group means and differences comparing the sum of log reaction times (seconds) on the Trails B task depending on data source (medical vs commercial), on-road test outcome (safe vs unsafe), and stage of the Trails B task (9 target vs 18 target). The Trails B 9 Target plot uses a y-axis break to improve readability of outliers. Vehicle drivers who failed the road test were slower on the Trails B task than drivers who passed the road test.

On the Modified TMT-A 9-target task, post-hoc tests did not confirm any statistical differences in error counts between participants from the medical and commercial data sources, whether they were classified as unsafe in the on-road assessment or classified as safe (Figure 13). Likewise, on the Modified TMT-A 18-target task, post-hoc tests did not confirm any statistical differences in error counts between participants from the medical and commercial data sources, whether they were classified as unsafe in the on-road assessment or classified as safe.

**Figure 13:**
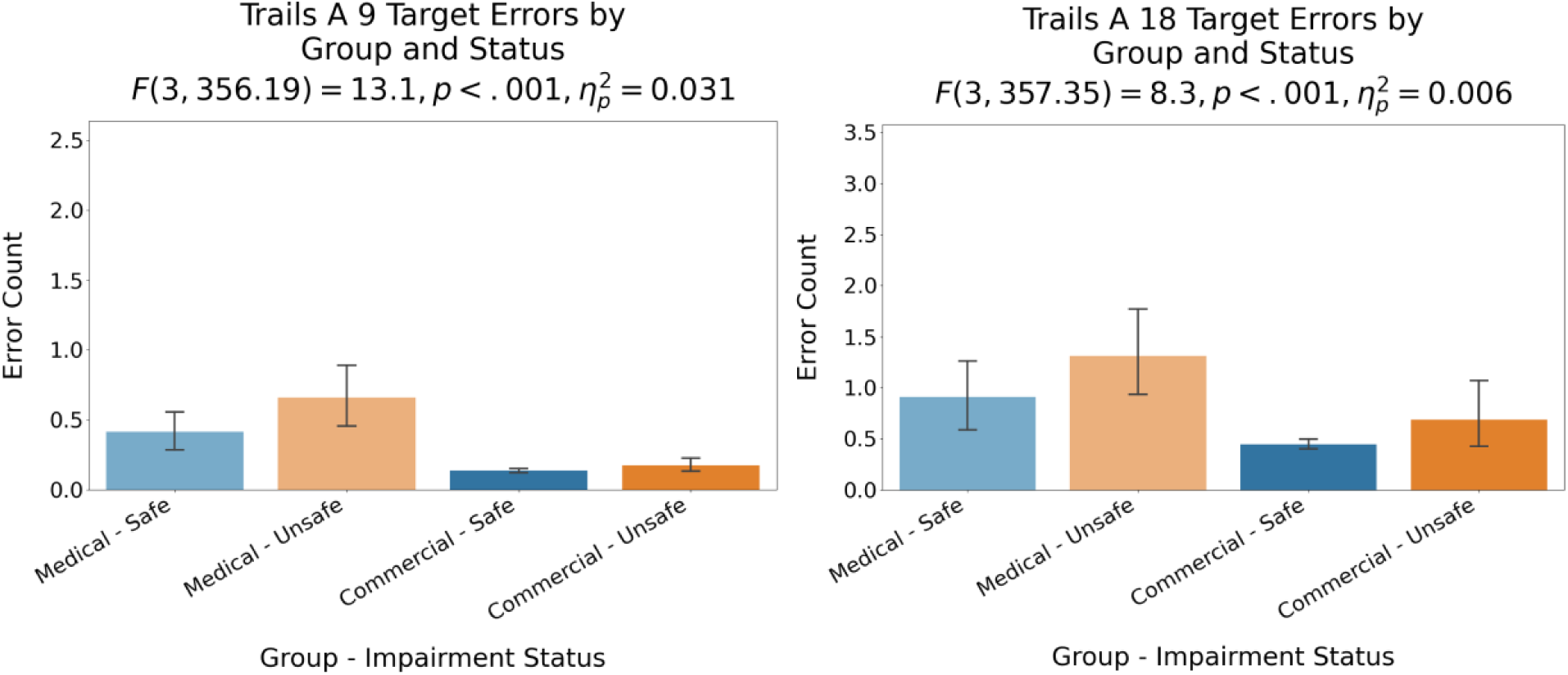
Group means and differences comparing total error count on the Trails A task depending on data source (medical vs commercial), on-road test outcome (safe vs unsafe), and stage of the Trails A task (9 target vs 18 target). Vehicle drivers who failed the road test accumulated more errors on the Trails A task than drivers who passed the road test.

On the Modified TMT-B 9-target task, post-hoc tests did not confirm any statistical differences in error counts between participants from the medical and commercial data sources, whether they were classified as unsafe in the on-road assessment or classified as safe (Figure 14). Likewise, on the Modified TMT-B 18-target task, post-hoc tests did not confirm any statistical differences in error counts between participants from the medical and commercial data sources, whether they were classified as unsafe in the on-road assessment or classified as safe.

**Figure 14:**
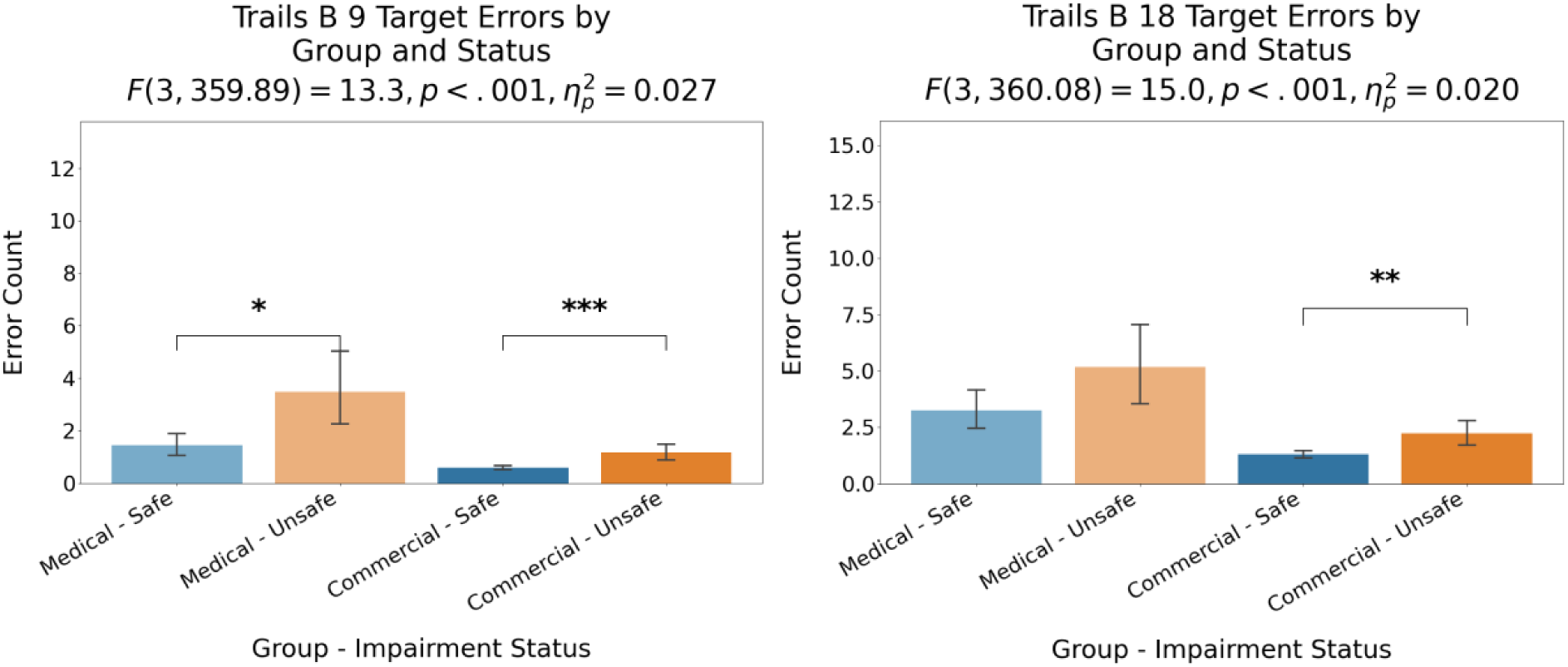
Group means and differences comparing total error count on the Trails B task depending on data source (medical vs commercial), on-road test outcome (safe vs unsafe), and stage of the Trails B task (9 target vs 18 target). Vehicle drivers who failed the road test accumulated more errors on the Trails B task than drivers who passed the road test.

On the Modified TMT-B 9-target task, participants from the medical data source had significantly higher error counts if they were classified as unsafe on the on-road assessment compared to if they were classified as safe (Figure 14). On the same 9-target task, participants from the commercial data source also had significantly higher error counts if they were classified as unsafe compared to if they were classified as safe. Likewise, on the Modified TMT-B 18-target task, commercial participants classified as unsafe made significantly more errors than those classified as safe. No such difference was observed in the medical data source for the 18-target configuration.

**Table 7:** Summary of one-way ANOVA results for performance metrics by safety group and data source (n=4719). For all variables, p<0.5 for Levene test, therefore Welch’s ANOVA was used instead of regular ANOVA.

| Metric | SS_Effect | MS_Effect | SS_Error | MS_Error | df | F | p | $\eta^2p$ |
| --- | --- | --- | --- | --- | --- | --- | --- | --- |
| Motor 9 Target Log RT | 4690.09 | 1563.36 | 20241.98 | 4.29 | 3, 361.01 | 154.64 | <.001 | .188 |
| Motor 18 Target Log RT | 11865.91 | 3955.30 | 60969.12 | 12.93 | 3, 366.02 | 185.89 | <.001 | .163 |
| Trails A 9 Target Log RT | 4808.31 | 1602.77 | 26268.28 | 5.57 | 3, 363.60 | 145.63 | <.001 | .155 |
| Trails A 18 Target Log RT | 24718.08 | 8239.36 | 143887.73 | 30.52 | 3, 360.82 | 154.04 | <.001 | .147 |
| Trails B 9 Target Log RT | 26008.31 | 8669.44 | 176135.94 | 37.36 | 3, 355.24 | 63.55 | <.001 | .129 |
| Trails B 18 Target Log RT | 103810.52 | 34603.51 | 398703.65 | 84.56 | 3, 356.65 | 113.48 | <.001 | .207 |
| Trails A 9 Target Errors | 47.74 | 15.91 | 1516.67 | 0.32 | 3, 356.19 | 13.11 | <.001 | .031 |
| Trails A 18 Target Errors | 151.44 | 50.48 | 23483.53 | 4.98 | 3, 357.35 | 8.25 | <.001 | .006 |
| Trails B 9 Target Errors | 1314.33 | 438.11 | 48255.90 | 10.23 | 3, 359.89 | 13.32 | <.001 | .027 |
| Trails B 18 Target Errors | 2836.47 | 945.49 | 142543.99 | 30.23 | 3, 360.08 | 15.02 | <.001 | .020 |

**Table 8:** T-test comparison of means by safety group in the full medical dataset (n=314, * = p < .05, † = p < .10, n.s. = not significant).

| Metric | N-safe | N-unsafe | N-total | P-Value | Significance |
| --- | --- | --- | --- | --- | --- |
| Motor 9 Target Log RT | 176 | 138 | 314 | 0.0001 | *** |
| Motor 18 Target Log RT | 176 | 138 | 314 | 0.0000 | *** |
| Trails A 9 Target Log RT | 176 | 138 | 314 | 0.0001 | *** |
| Trails A 18 Target Log RT | 176 | 138 | 314 | 0.0001 | *** |
| Trails B 9 Target Log RT | 176 | 138 | 314 | 0.0019 | ** |
| Trails B 18 Target Log RT | 176 | 138 | 314 | 0.0008 | *** |
| Trails A 9 Target Errors | 176 | 138 | 314 | 0.0611 | - |
| Trails A 18 Target Errors | 176 | 138 | 314 | 0.1393 | - |
| Trails B 9 Target Errors | 176 | 138 | 314 | 0.0116 | * |
| Trails B 18 Target Errors | 176 | 138 | 314 | 0.0546 | - |

**Table 9:** T-test comparison of means by safety group in the full commercial dataset (n=4405).

| Metric | N-safe | N-unsafe | N-total | P-Value | Significance |
| --- | --- | --- | --- | --- | --- |
| Motor 9 Target Log RT | 3733 | 672 | 4405 | 0.0000 | *** |
| Motor 18 Target Log RT | 3733 | 672 | 4405 | 0.0002 | *** |
| Trails A 9 Target Log RT | 3733 | 672 | 4405 | 0.0000 | *** |
| Trails A 18 Target Log RT | 3733 | 672 | 4405 | 0.0000 | *** |
| Trails B 9 Target Log RT | 3733 | 672 | 4405 | 0.0000 | *** |
| Trails B 18 Target Log RT | 3733 | 672 | 4405 | 0.0000 | *** |
| Trails A 9 Target Errors | 3733 | 672 | 4405 | 0.1404 | - |
| Trails A 18 Target Errors | 3733 | 672 | 4405 | 0.1556 | - |
| Trails B 9 Target Errors | 3733 | 672 | 4405 | 0.0005 | *** |
| Trails B 18 Target Errors | 3733 | 672 | 4405 | 0.0014 | ** |

### 3.3 Model Evaluation

#### 3.3.1 Unstratified model

All unstratified data fit on the logistic regression models had significantly converged. The medical data group model had a significant likelihood ratio test (p<0.001) (Table 10), the commercial data group model had a significant likelihood ratio test (p<0.001) (Table 11), and the combined data group model had a significant likelihood ratio test (p<0.001) (Table 12). The significant likelihood ratio test and McFadden Pseudo-R² demonstrate that the model is able to improve predictive power over random chance (Hosmer & Lemeshow, 2013; Seabold & Perktold, 2010). There were small differences between area under the curve (AUC) metrics between the training and test splits when evaluating the unstratified model fit on medical, commercial, and combined data sources (Figure 15). The tests were performed using the statsmodel python library and common standard when quantifying the model fit of a Logistic Regression model (Seabold & Perktold, 2010). When the models were evaluated against the testing dataset, there were near monotonic increasing relationships between the Neurapulse tasks and the probability of failing the on-road test.

**Table 10:** Model fit tables for the medical unstratified training dataset (* = p < .05, † = p < .10, n.s. = not significant)

| Term | Value |
| --- | --- |
| N (observations) | 157 |
| Df (model) | 3 |
| Log-likelihood | -102.54 |
| Pseudo-$R^2$ (McFadden) | 0.056 |
| LLR p-value | 0.0066 |
| Converged? | Yes |

| Predictor | Sign | Significance |
| --- | --- | --- |
| Intercept | – | * |
| Motor 18 Target (Log RT) | + | † |
| Trails A 18 Target (Log RT) | + | n.s. |
| Trails B 18 Target (Log RT) | + | n.s. |

**Table 11:** Model fit tables for the commercial unstratified training dataset (* = p < .05, n.s. = not significant)

| Term | Value |
| --- | --- |
| N (observations) | 2203 |
| Df (model) | 3 |
| Log-likelihood | -916.92 |
| Pseudo-$R^2$ (McFadden) | 0.025 |
| LLR p-value | 2.39e-10 |
| Converged? | Yes |

| Predictor | Sign | Significance |
| --- | --- | --- |
| Intercept | – | * |
| Motor 18 Target (Log RT) | + | n.s. |
| Trails A 18 Target (Log RT) | + | * |
| Trails B 18 Target (Log RT) | + | * |

**Table 12:** Model fit tables for the combined (medical and commercial) unstratified training dataset (* = p < .05, n.s. = not significant)

| Term | Value |
| --- | --- |
| N (observations) | 2360 |
| Df (model) | 3 |
| Log-likelihood | -1027.8 |
| Pseudo-$R^2$ (McFadden) | 0.058 |
| LLR p-value | 2.54e-27 |
| Converged? | Yes |

| Predictor | Sign | Significance |
| --- | --- | --- |
| Intercept | – | * |
| Motor 18 Target (Log RT) | + | * |
| Trails A 18 Target (Log RT) | + | * |
| Trails B 18 Target (Log RT) | + | * |

**Table 13:** Training and test split counts for medical, commercial, and combined data sources. The class distribution is not highly skewed.

| Group | Subset | N | Safe | Unsafe |
| --- | --- | --- | --- | --- |
| Medical | Training | 157 | 82 | 75 |
| Medical | Test | 157 | 94 | 63 |
| Commercial | Training | 2203 | 1867 | 336 |
| Commercial | Test | 2202 | 1866 | 336 |
| All | Training | 2360 | 1949 | 411 |
| All | Test | 2359 | 1960 | 399 |

**Figure 15:**
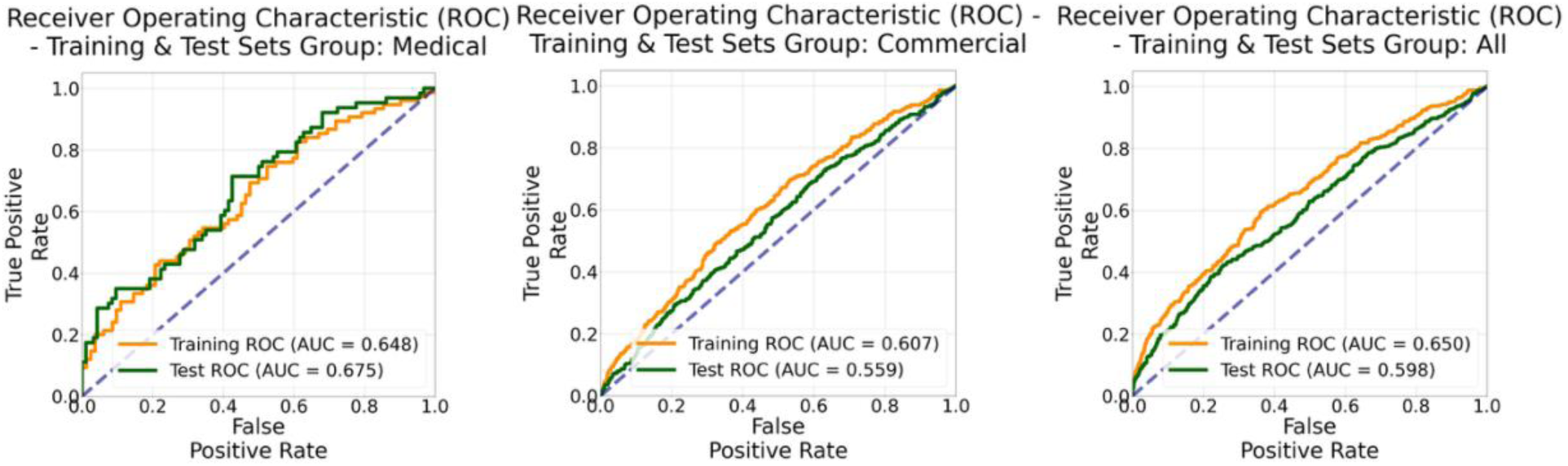
ROC Curve plots for medical, commercial, and combined data sources, respectively. Differences in AUC were smallest in the medical data source.

**Figure 16:**
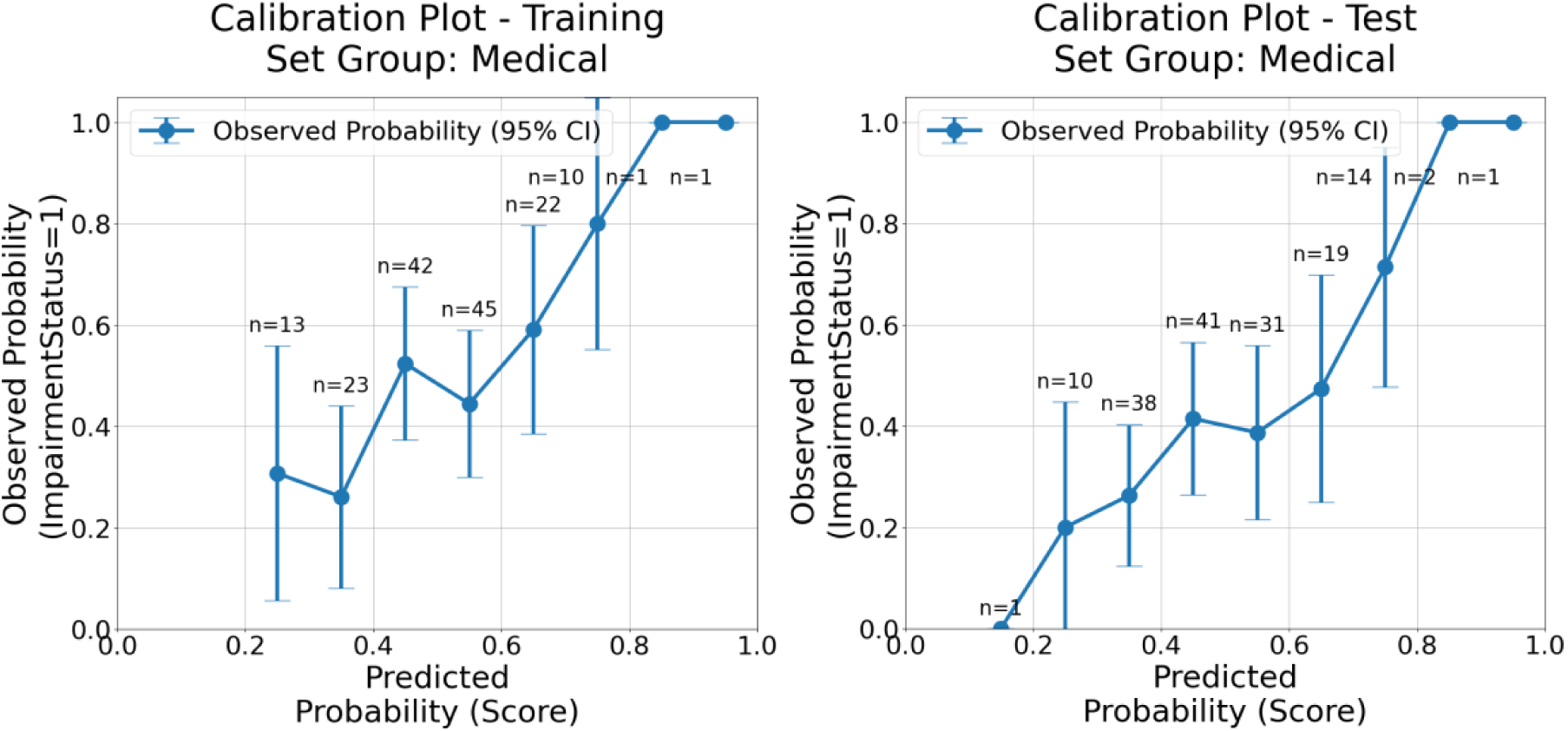
Calibration plot comparing predicted probability of on-road failure using the Motor/Trails task battery logistic model vs the actual probability of an unsafe on-road test outcome. Error bars represent 95% CI levels. Both unstratified medical training (left) and test data (right) groups showed a near monotonic increasing relationship.

**Figure 17:**
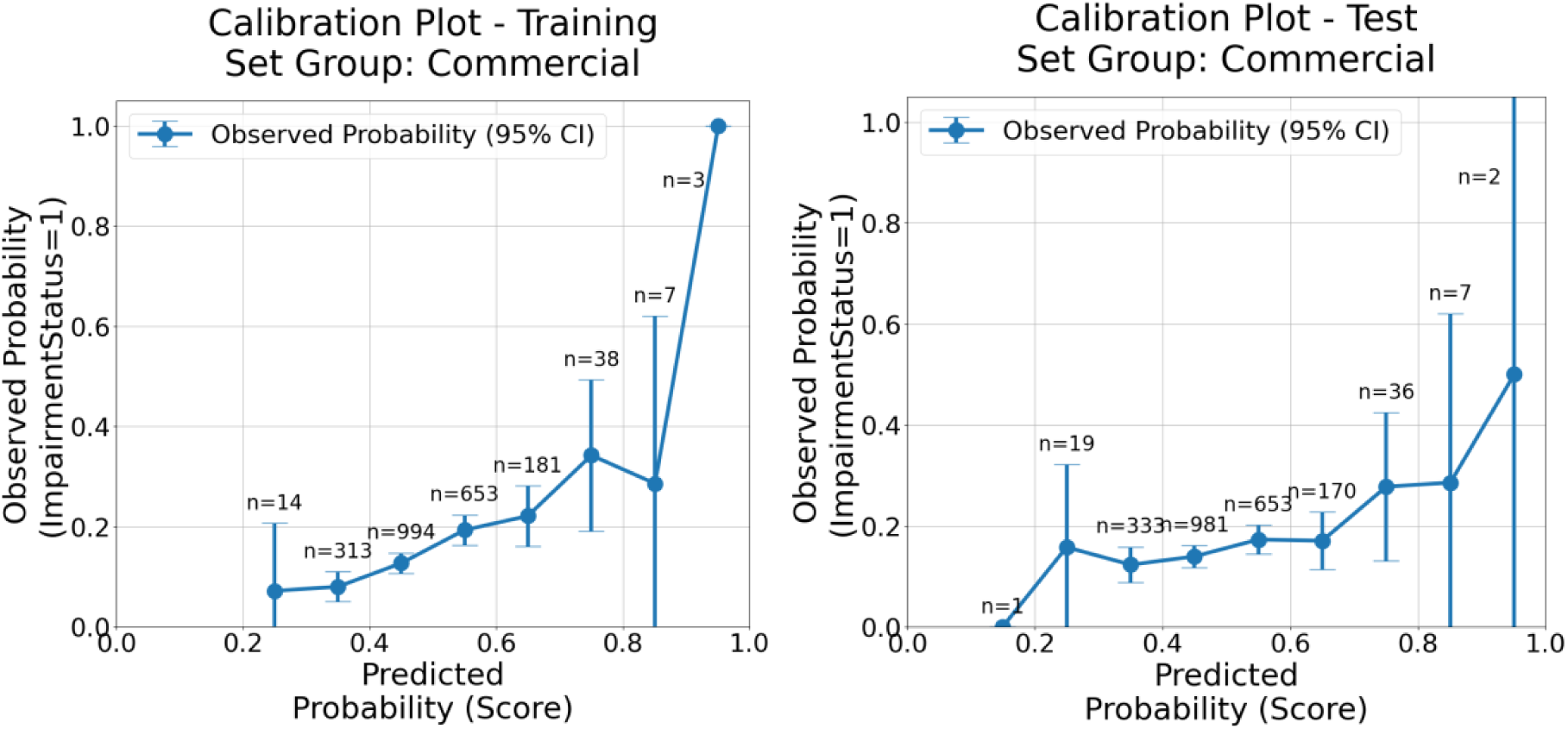
Calibration plot comparing predicted probability of on-road failure using the Motor/Trails task battery logistic model vs the actual probability of an unsafe on-road test outcome. Error bars represent 95% CI levels. Both unstratified commercial training (left) and test data (right) groups showed a near monotonic increasing relationship.

**Figure 18:**
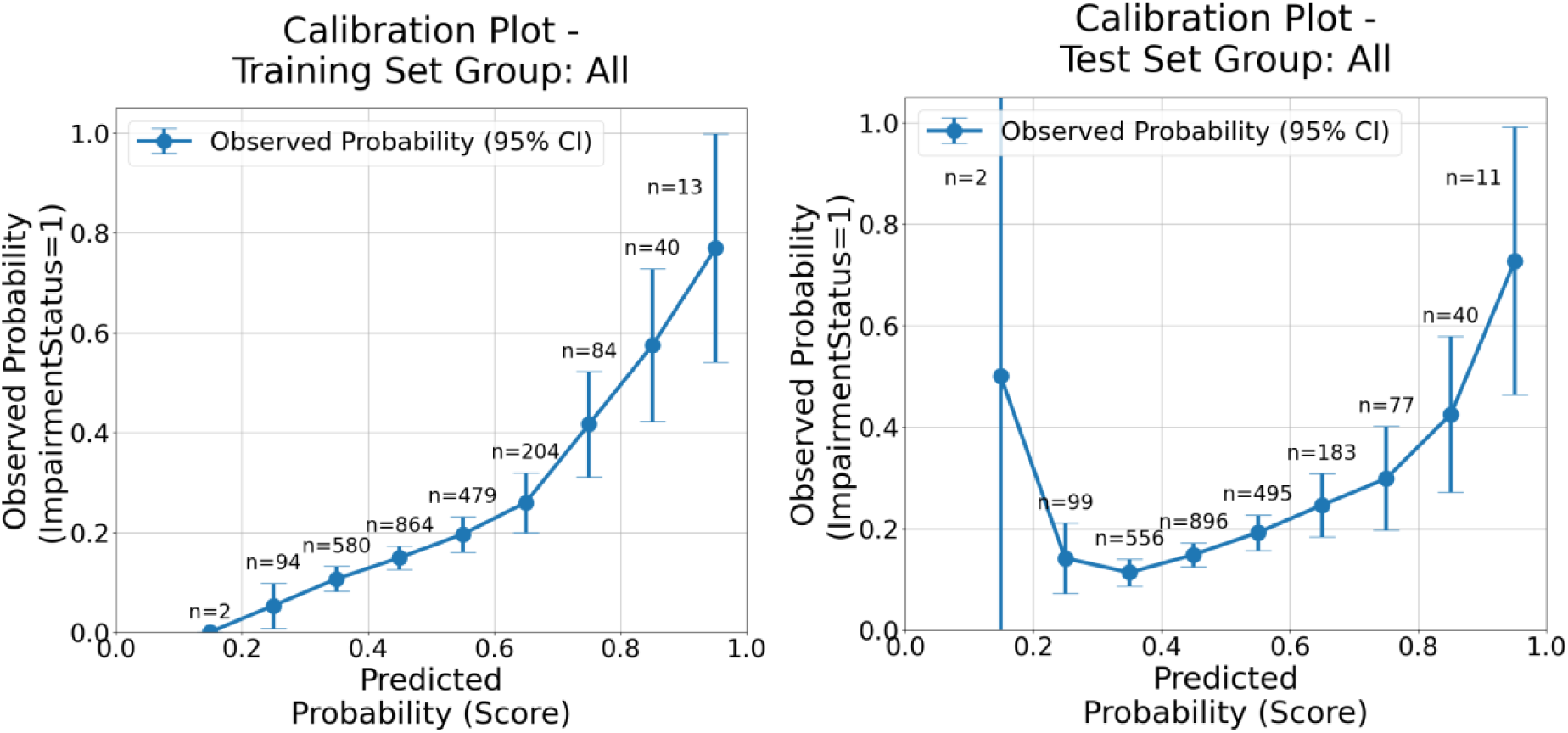
Calibration plot comparing predicted probability of on-road failure using the Motor/Trails task battery logistic model vs the actual probability of an unsafe on-road test outcome. Error bars represent 95% CI levels. Both unstratified training (left) and test data (right) groups from the combined full data source showed a near monotonic increasing relationship.

#### 3.3.2 Updated Accuracy Metrics After Stratification

**Table 14:** Training and test split counts for medical, commercial, and combined data sources. The class distribution is not highly skewed. Since the test set is held out, it is not stratified.

| Group | Subset | N | Safe | Unsafe |
| --- | --- | --- | --- | --- |
| Medical | Training | 3734 | 1867 | 1867 |
| Medical | Test | 157 | 94 | 63 |
| Commercial | Training | 3734 | 1867 | 1867 |
| Commercial | Test | 2202 | 1866 | 336 |
| All | Training | 7468 | 3734 | 3734 |
| All | Test | 2359 | 1960 | 399 |

There were small differences between AUC metrics between the training and test splits when evaluating the stratified model fit on medical and commercial. The differences substantially decreased in the combined model after stratification.

**Figure 19:**
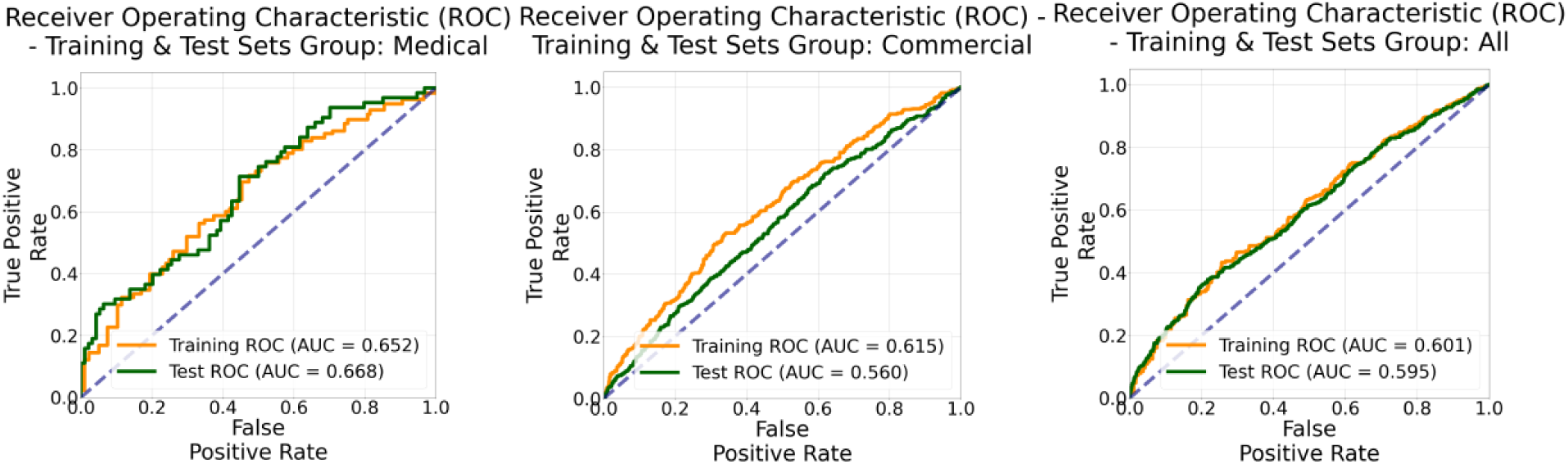
ROC Curve plots for medical, commercial, and combined data sources, respectively. Differences in AUC were smallest in the combined data source.

**Figure 20:**
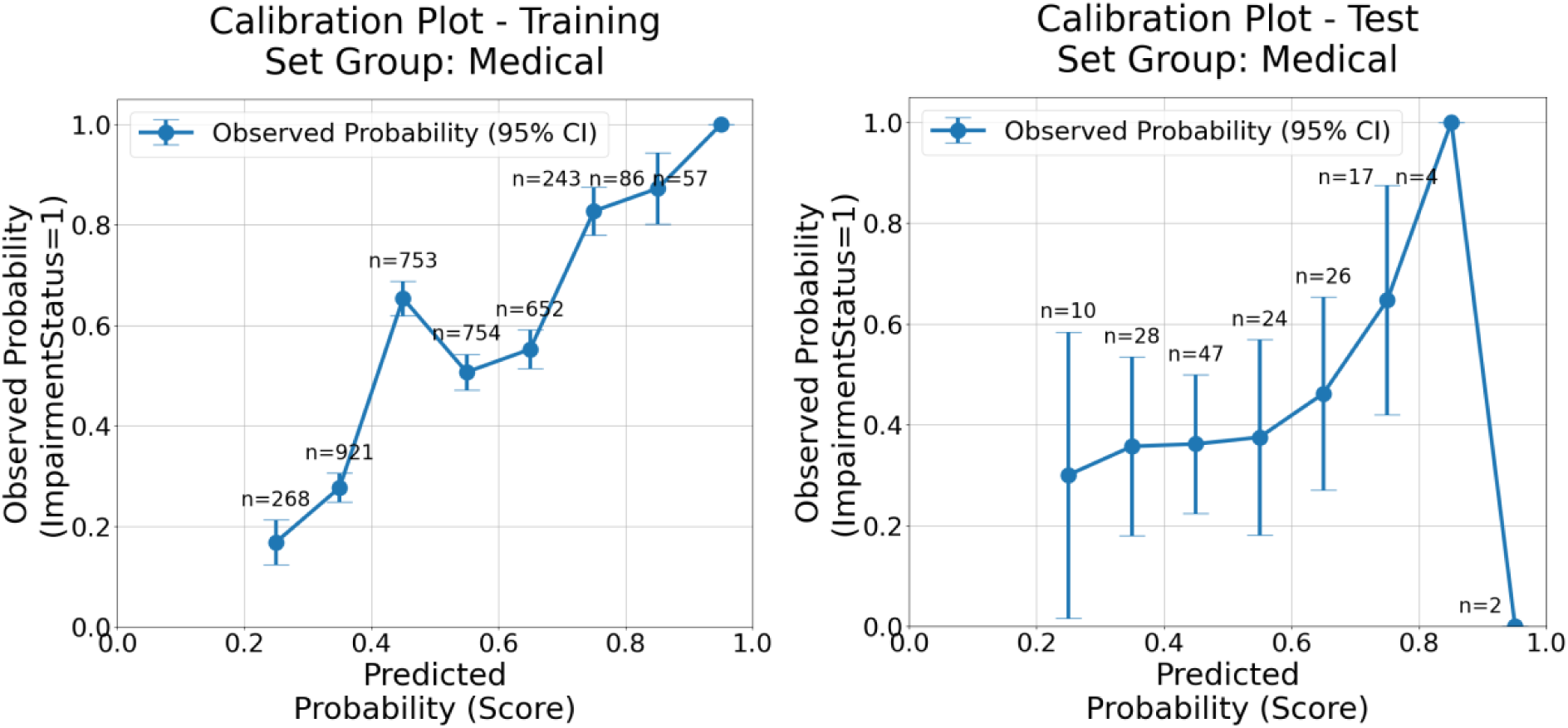
Calibration plot comparing predicted probability of on-road failure using the Motor/Trails task battery logistic model vs the actual probability of an unsafe on-road test outcome. Error bars represent 95% CI levels. Both stratified medical training (left) and unstratified test data (right) groups showed a near monotonic increasing relationship.

**Figure 21:**
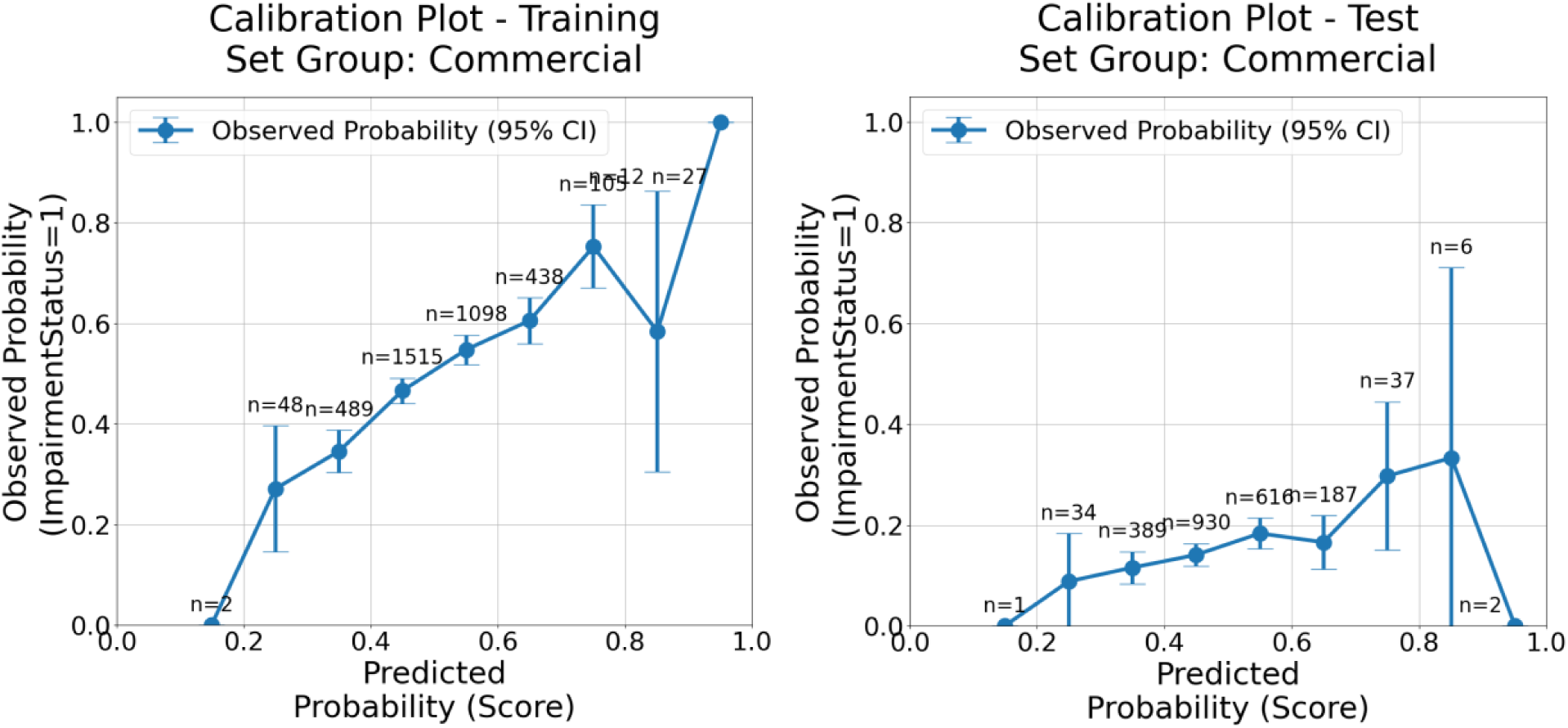
Calibration plot comparing predicted probability of on-road failure using the Motor/Trails task battery logistic model vs the actual probability of an unsafe on-road test outcome. Error bars represent 95% CI levels. Both stratified commercial training (left) and unstratified test data (right) groups showed a near monotonic increasing relationship.

**Figure 22:**
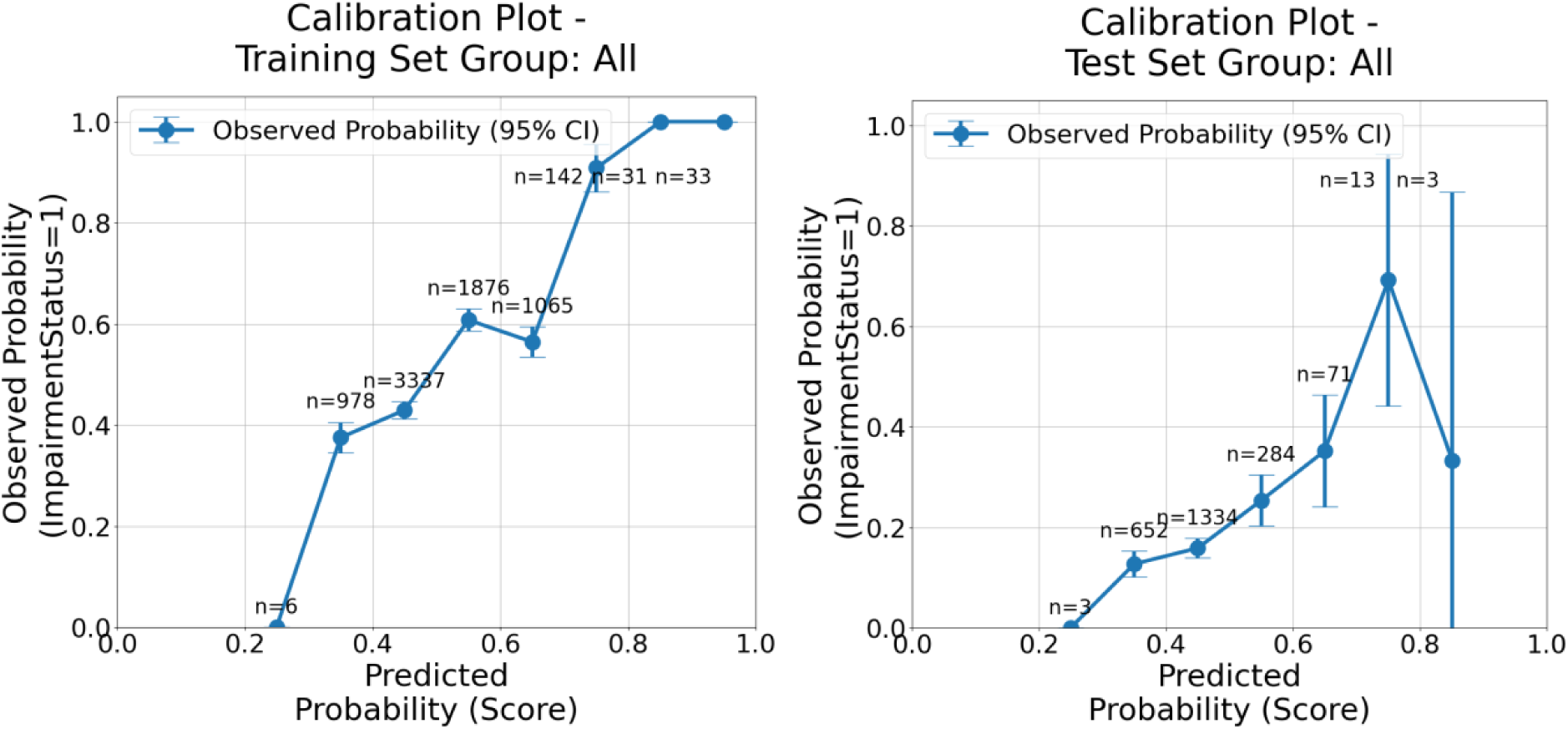
Calibration plot comparing predicted probability of on-road failure using the Motor/Trails task battery logistic model vs the actual probability of an unsafe on-road test outcome. Error bars represent 95% CI levels. Both stratified training (left) and unstratified test data (right) groups from the combined full data source showed a near monotonic increasing relationship.

#### 3.3.3 Detailed Accuracy Metrics

Using the methodology from Scott et al. (2023) for setting an optimal threshold, we used the function objective= FP*FP_Weight + FN*FN_Weight, where FP is the count of false positives in the training dataset, FN is the count of false negatives in the training dataset, and FP_Weight and FN_Weight are a custom weight to represent the cost of error for FP and FN cases, respectively. The penalties quantify negative consequences across dimensions of participant mental hazard (false positive) and driving accident cost (false negative) (Scott et al., 2023).

Using a single dichotomous threshold, the maximum objective score was selected at 0.615 (Figure 23). The stratified logistic regression model trained on the combined dataset was used to generate a predicted probability for each participant in the held-out test set. For each case, if the predicated probability was greater than or equal to the cutpoint threshold, they were classified as “unsafe”, otherwise they were classified as “safe”. The threshold of 0.615 prioritized a low sensitivity and high specificity, which calibrated the results to minimize false positives (FP=61) while maximizing true negatives (TN=1899). Evaluating the testing set from the combined data group resulted in a positive predictive value (PPV) of 43.0%, a negative predictive value (NPV) of 84.3%, and an accuracy (ACC) of 82.5%. Evaluating the testing set from the commercial data group resulted in a positive predictive value (PPV) of 34.4%, a negative predictive value (NPV) of 85.3%, and an accuracy (ACC) of 83.8%. Evaluating the testing set from the medical data group resulted in a positive predictive value (PPV) of 55.8%, a negative predictive value (NPV) of 65.8%, and an accuracy (ACC) of 63.1%. When comparing like-for-like commercial data groups between the previous Vitals tasks and our new tasks, the logistic model shows similar performance and an improved classification rate. The Scott et al. (2023) and Atkin et al. (2024) method of using a trichotomous cutpoint excluded a proportion of participants from the accuracy metrics. In our new dichotomous cutpoint regime, every participant’s results are included in the accuracy metrics.

**Figure 23:**
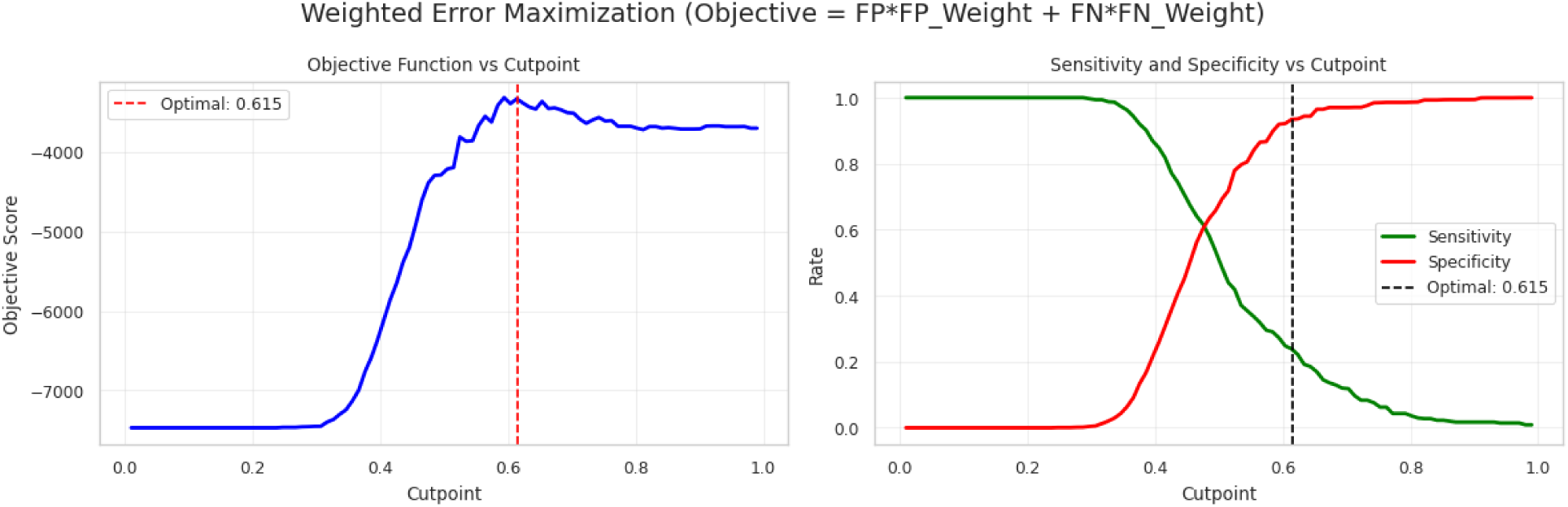
Objective function plot for model cutpoint using FP*FP_Weight + FN*FN_Weight formula. A cutpoint of 0.615 maximizes the objective score.

**Table 15.** Accuracy metrics for the held-out test set (all data sources) generated with the single dichotomous cut-point selected by the FP/FN-weighted optimization method. In the like-for-like commercial data groups, our new task model achieved higher accuracy and negative predictive value, but lower positive predictive value, compared to the previous task model reported by Scott et al. (2023) and Atkin et al. (2024).

| Data Source | Decision rule | TN | TP | FN | FP | NPV | PPV | ACC | N |
| --- | --- | --- | --- | --- | --- | --- | --- | --- | --- |
| New Tasks – All sources | 1 cut-point 0.615 | 1899 | 46 | 353 | 61 | 84.3% | 43.0% | 82.5% | 2359 |
| New Tasks – Medical | 1 cut-point 0.615 | 75 | 24 | 39 | 19 | 65.8% | 55.8% | 63.1% | 157 |
| New Tasks – Commercial | 1 cut-point 0.615 | 1824 | 22 | 314 | 42 | 85.3% | 34.4% | 83.8% | 2202 |
| Previous Tasks - Commercial (Scott et al., 2023) | 2 cut-points 0.48/0.68 | 1330 | 41 | 293 | 60 | 81.9% | 40.6% | 79.5% | 1724 |
| Previous Tasks - Commercial (Atkin et al., 2024) | 2 cut-points 0.48/0.68 | 1490 | 37 | 337 | 60 | 81.6% | 38.0% | 79.4% | 1924 |

## 4. Discussion and Limitations

In this study, we examined performance on the Neurapulse cognitive assessment tool in mostly older, medically at-risk drivers and commercial drivers. Drivers also completed an on-road evaluation of their driving abilities, and a logistic regression model was used to predict the outcome of this evaluation based on Neurapulse performance. Neurapulse consists of three tasks – a motor task, and the Trail–Making Tests A and B – and can be completed in approximately 5 minutes using a smartphone. This represents a significant advancement in administration efficiency compared to a previous and effective assessment tool called Vitals, which has a completion time of approximately 30 minutes and must be performed in-office using a tablet (Atkin et al., 2024; Bakhtiari et al., 2020; Scott et al., 2023). More importantly, Neurapulse achieved slightly higher classification accuracy in commercial drivers compared to Vitals, with higher negative predictive value but lower positive predictive value. In the medically at-risk drivers, overall classification accuracy was lower than for commercial drivers, but with a higher positive predictive value, allowing for greater confidence that at-risk drivers who fail the Neurapulse tasks are likely to fail an on-road evaluation. Because it achieves improved classification accuracy despite a dramatically reduced test completion time and increased accessibility, the Neurapulse assessment tool demonstrates incremental validity in predictive power and practical utility over the previous Vitals tool.

In older or medically at-risk drivers, current research largely supports the utility of the TMT for predicting on-road safety (Asimakopulos et al., 2012; CMA, 2025; Costello et al., 2024; Duncanson et al., 2018; Rashid et al., 2020). With a 63.1% classification accuracy, our study also supports this conclusion. There are a few negative studies on this question however, most notably Vaucher et al. (2014). This study found that older drivers who failed the TMT were much more likely to fail a driving evaluation, but nevertheless concluded that the TMT lacks sufficient sensitivity, specificity, and PPV to recommend driving cessation on this basis. While our older, at-risk sample had a lower NPV than participants in that study (65.8% vs. 96.9%), our PPV was much higher (55.8% vs. 9.5%). Vaucher et al. (2014) use a 150s cutoff score for Trails B that is lower than typical for this test, which would have lowered their false negative rate but increased their false positive rate. In our study we did not use cut-off scores, and participants were allowed to continue the test until completion, similar to Classen et al. (2013). Instead, we classified our drivers using a cutpoint derived from our logistic regression model, which used reaction times on the 18-dot motor task, 18-dot TMT-A, and 18-dot TMT-B as predictor variables. Drivers were classified as unsafe if the model assigned them a 61.5% probability or higher of failing the on-road evaluation. Using this approach, we were able to achieve a much higher PPV than Vaucher et al. (2014), with an AUC that is very similar to their results.

Increasing the cutpoint could improve our model’s PPV further, but at the expense of lower NPV. Overall, Neurapulse appears to have some value in predicting on-road driving performance in medically at-risk drivers, and its accessibility and short administration time makes it both more convenient and more easily incorporated into a battery of other assessments relative to other versions of the TMT. In terms of univariate statistics, drivers who passed or failed the on-road evaluation differed significantly on numerous measures of the Neurapulse tasks. Specifically, drivers in both the medically at-risk and commercial driving groups differed on every measure of reaction time, including both the 9-target and 18-target levels of each task. By contrast, commercial drivers only differed in terms of error rate on both levels of the TMT-B task, while older, at-risk drivers only differed on the 9-target level of TMT-B. This is consistent with past TMT research, which typically reveals a relationship between driving performance and completion time (Ball et al., 2006; Carr et al., 2011; Classen et al., 2008, 2013, 2015; Friedman et al., 2013; Krasniuk et al., 2023; Marshall et al., 2007; Motta et al., 2024; Ott et al., 2013; Stutts et al., 1998; Tarawneh et al., 1993). Error rate less often assessed, although it is known to be associated with MCI and dementia (Ashendorf et al., 2008). Ott et al. (2013) found significant differences in TMT-B error rate between safe and unsafe older drivers, some of whom had been diagnosed with cognitive impairment, while Venkatesan et al. (2018) found that TMT-B error rate was associated with specific risky driving behaviours in early dementia patients. Our results likewise show an association between TMT-B error rate and risky on-road driving. The similarity between our results and these previous studies suggests that Neurapulse can be used instead of desktop or pen-and-paper versions of the TMT, allowing tests to be administered remotely regardless of time and location.

Despite its short administration time relative to the Vitals task battery, Neurapulse tests a similar range of cognitive and sensorimotor functions, including many of the functions considered essential for driving (CCMTA 2022). Compared to the Vitals tasks, however, which were heavily modified from various neuropsychological assessments (Atkin, 2025; Bakhtiari et al., 2020) the TMT is a well-established neuropsychological assessment, with decades of research showing that TMT-A tests visual search and processing speed, while TMT-B tests executive functions such as cognitive flexibility and divided attention. Neurapulse uses a lightly-modified version of the TMT, with two sequences of numbers for TMT-B rather than one sequence of numbers and one sequence of letters, and an algorithmically-generated dot configuration pattern. We can therefore be more confident with Neurapulse that unsafe drivers are unsafe because of impairment to functions that are important for driving. However, we are unable to diagnose the reasons for this impairment. Cognitive impairment is associated with many neurological conditions and mental states, each with their own etiology and prognosis for recovery (Hancock & Desmond, 2000; Saxby et al., 2013). We also cannot say whether each driver’s poor performance is transient or persistent, as their medical status is unknown and performance was only measured once. Further observation and testing of unsafe drivers would be required to identify the source of their problem.

### 4.1 Limitations

This study found an association between TMT-B error rate and unsafe driving in commercial drivers, but not in medically at-risk drivers. Safe commercial drivers rarely committed errors (Figure 14), so even a small increase in errors amongst unsafe drivers would be statistically and practically significant. By contrast, even safe medical drivers committed multiple errors on average, more than the unsafe commercial drivers, so there was less distinction between safe and unsafe older drivers in terms of their error rate. The sample size for the medically at-risk driving group is an order of magnitude smaller than for the commercial drivers, however, with less power to detect differences, so it is also possible that a difference in error rate would be observed in medically at-risk drivers if a larger sample had been tested.

Our two driving groups differed not just in average age, but also in their distribution of ages. For the medical group, we had limited age bins at the lower range, while for the commercial group, we had limited age bins at the upper range. This resulted in lower coherence in our age-stratified model when compared to the unstratified model. However, stratification did produce a model with a more stable AUC, suggesting that the predictive power in the unstratified model may be overstated due to the random sampling procedure in the training sample. For this reason, the stratified model should be preferred over the unstratified model to maximize generalization with new samples.

For TMT-B, we modified the task to alternate between two numerical sequences (1-25-2-26-) rather than between a numerical and an alphabetic sequence (1-A-2-B-). This was done to minimize cultural bias on the task, which might otherwise disadvantage test-takers whose first language does not use the Latin alphabet (Bezdicek et al., 2016, Zhao et al., 2013). Some languages use a different numerical system than Arabic numerals, and the TMT has occasionally been modified to use alternative numerical systems (Cova et al., 2021). Thus, there could still be cultural and linguistic biases in our study. However, we expect that this bias would be less than for the original TMT-B task, due to the widespread use of Arabic numerals even in languages that do not use the Latin alphabet. A future study could test this expectation by using Bezdicek et al.’s (2016) methodology to compare the original TMT-B task with our modified version.

As with previous studies from our group, gender and other demographic information other than age was not collected (Atkin et al., 2024; Scott et al., 2023).

### 4.2 Conclusion

Rapid assessment of cognitive abilities has the potential to improve driver safety while minimizing burden to test-takers and administrators. The results of this study suggest that a rapid and accessible version of the TMT can predict unsafe driving with comparable accuracy to more time-consuming and administratively burdensome means of testing.

## Acknowledgements

The authors would like to thank Dr. Craig Chapman and Quinn Boser for their support in algorithmic development for the trails task dot generation, as well as task selection.

